# A protein-fragment complementation assay to quantify synthetic protein scaffold efficiency

**DOI:** 10.1101/2025.03.19.640584

**Authors:** Pascale Lemieux, Alexandre K Dubé, Christian R Landry

## Abstract

Scaffolds are powerful tools in synthetic biology used for various applications, from increasing yield to optimizing signalling specificity. Protein scaffolds can be built by fusing peptide binding domains (PBD) and attaching the peptide they bind to the enzymes, inducing spatial proximity. Only a few PBD-peptide combinations have been tested in this context, and no combination produced a high yield in yeast, an important chassis in biotechnology. Therefore, there is a need for more exploration of PBD-peptide pairs to be used in this model. Scaffold characterization is challenging because it is often dependent on a model pathway with an output that is difficult to measure quantitatively. Here, we use a protein-fragment complementation assay (PCA) to study scaffolding efficiency in yeast, which allows to couple scaffolding efficiency with growth rate. First, we characterize the strength of PBD-peptide interactions (PPI) and the binding availability of the PBDs and peptides. Then, we test different scaffold architectures and expression levels to quantify the simultaneous binding of peptide pairs to the scaffold. We show that PPI strength of the weakest binding PBD-peptide pair is critical for scaffolding efficiency and that PPI strength is limited by low binding availability of some domains and peptides *in vivo*. Also, we find that slight architectural variations and expression levels have a significant impact on scaffolding efficiency detected by DHFR PCA. Finally, we used DHFR PCA approaches to characterize novel PBD-peptide pairs and we identified pairs to expand the sequence toolbox for scaffold design in yeast through DHFR PCA easy-to-read signal.

## Introduction

One challenge in synthetic biology is the optimization of the efficiency and specificity of synthetic pathways^1^. In nature, we often observe highly efficient pathways in which proteins operate in close proximity, thus accelerating signal transduction or inducing metabolic channeling. Such endogenous pathways involve the colocalization of enzymes by direct physical interactions^2,3^ or via a scaffold molecule^4–6^, creating large protein complexes. In such cases, protein stoichiometry and interaction specificity contribute to maintaining function and correct protein complex assembly^7,8^. The regulation of spatial proximity is also important as it prevents intermediary metabolites from diffusing in the cytoplasm, which can slow down pathways^9,10^. The multiple layers of control and the spatial organisation we observe for endogenous pathways are absent when we insert a synthetic pathway in a host cell.

Since the proteins of synthetic pathways are often of exogenous origin, they do not have a constrained localization in the cell encoded by their sequence or by specific interactions with endogenous proteins. Therefore, the enzymes of a synthetic metabolic pathway can be scattered in the cytoplasm and thus depend on the diffusion of intermediary metabolites to produce the end molecule. The crowding of the cellular environment could slow down intermediary metabolite diffusion between the enzymes of the synthetic pathway even more, limiting its yield^11^. Intermediary metabolite diffusion is further slowed down due to the lack of proximity with the enzyme catalysing the subsequent reaction of the pathway^12,13^. Moreover, many pathways produce toxic metabolites that impair the growth of the host cell when they are allowed to diffuse. One approach tested to compensate for the low yield of synthetic pathways was to over-express the exogenous enzymes^14^, which was found to hinder cell growth by increasing the metabolic burden from the expression of the synthetic pathway^12,15,16^. Inspired by what is found in nature, another solution is the use of a synthetic scaffold, which artificially induces the proximity of enzymes of a pathway to increase its efficiency^17^.

Design features such as the modification of enzyme sequences, which could hinder their activity or folding, and the modular control on the recruitment of specific enzymes are important for synthetic scaffold design. One possibility to limit altering enzyme activity and ensure specificity is to construct artificial scaffolds by fusing peptide binding domains (PBD)^17^. Short peptides (∼10 a.a.) recognized by the PBDs can be fused to the enzymes of the pathway, which will result in bringing them together in the presence of the scaffold. The short peptides are expected to have a weaker impact on protein function in comparison to large fusion proteins^1^, and the intrinsic specificity of the PBDs enables the binding of their favored peptide fused to enzymes of the pathway, ensuring modularity^18,19^.

Scaffolding efficiency, which is mainly determined by the colocalization of enzymes of the synthetic pathway, is achieved when the PBDs simultaneously bind the peptides fused to the enzymes of interest. Therefore, scaffolding efficiency is dependent on the binding strength between the PBD-peptide pairs used in the scaffold. In *Escherichia coli*, this scaffolding strategy was shown to increase the efficiency of the heterologous mevalonate biosynthesis pathway by 77-fold and had success with other synthetic pathways^17,20–22^. However, the same approach showed a maximal yield increase of 5-fold in *Saccharomyces cerevisiae* (resveratrol biosynthesis), an important model for eukaryotic chassis in synthetic biology^23^. It is hard to identify a better scaffold sequence when very few PBD-peptide pairs have been tested *in vivo* for this application, and it remains challenging to characterize scaffolding efficiency without an easy-to-read output^24^. Previous studies generally used a heterologous pathway to characterize the increase in product yield, which is a proxy for pathway efficiency. However, specific metabolite purification methods are often required, and the findings may not be transferable to other pathways^23^. Indeed, the colocalization between the scaffold components is not directly characterized by this strategy. It rather assesses the effect of the scaffold on a specific pathway efficiency. Thus, there is a need for a method that would allow for direct quantification of the ability of a scaffold to bring components together (scaffolding efficiency) without relying on pathway efficiency as a readout.

The choice of PBDs and peptides contributing to the scaffold is crucial for its efficiency. The strength of the PBD-peptide interaction (PPI) *in vivo* is a determinant for the enzymes’ colocalization. Additionally, these heterologous proteins may also be degraded^25^ or interact with the proteome of the cell host in unexpected ways^26^. Both of these mechanisms reduce the amount of protein able to effectively bind other proteins in the cytosol and could interfere with the scaffold ability of colocalizing the enzymes. We refer to the free amount of protein or peptide able to bind as binding availability. An in-depth characterization of PBD-peptide scaffolds *in vivo*, while taking into account binding strength and binding availability of each PBD-peptide pair, is thus crucial for the development and exploration of more synthetic scaffolds, particularly in species used in synthetic biology applications such as yeast.

Here, we present an approach to measure scaffolding efficiency in *S. cerevisiae* with an easy-to-read output. It uses the dihydrofolate reductase protein-fragment complementation assay (DHFR PCA), which is a system with a growth-based read-out that is broadly used for protein-protein interaction quantification^19,27–29^. We use this reporter to characterize scaffolds built on two PBD-peptide pairs. Using heterologous PBD-peptide pairs described previously *in vitro*, we assayed their PPI strength and their binding availability *in vivo* with DHFR PCA^30^. We found that protein binding availability can profoundly impact PPI strength estimated *in vivo*. Then, we used DHFR PCA to assess the scaffolding efficiency for scaffolds designed with weak and strong binding PBD-peptide pairs. We also tested different scaffold architectures and examined their expression levels. We found that PPI strength, scaffold architecture and scaffold expression levels significantly impact the scaffolding efficiency. Our findings demonstrate that the DHFR PCA approach can capture variations in scaffolding efficiency caused by small differences in sequence and scaffold architecture. Finally, we individually tested the PPI strength and binding availability of more than 50 PBD-peptide pairs, which were not previously used in synthetic scaffolds. We identified candidate PBD-peptide pairs to add to the toolbox for scaffold design as a new resource for the community. In the future, DHFR PCA approaches could be applied to a diverse library of PBD-peptide to expand further the toolbox of sequences for scaffold design in yeast.

## Results

### PPI strength *in vivo* generally agrees with affinity values

We searched the literature for *in vitro* characterized PBD-peptide pairs with a wide range of affinity to understand how affinity impacts scaffolding efficiency. We were primarily interested in PBDs previously used in synthetic scaffolds, for which the affinity to one or multiple peptides had been characterized. We selected PBD sequences that include a SH3, a PDZ, and a GBD domain, which were shown to increase yield of synthetic pathways in bacteria and in yeast^17,20,23^. We also added the well-characterized C-terminal SH3 domain of GRB2p, referred to as GRB2^31,32^. We use capital letter abbreviations to name PBD sequences and lower-case abbreviations to name their binding peptides. We selected a binding peptide for the GBD and GRB2 domain, named respectively gbd1 and gab2^31,33^. For the SH3 and PDZ domains, multiple peptides were selected and numbered from the highest to lowest affinity as previously characterized *in vitro*, i.e. sh3.1 is the strongest binder tested, and sh3.4 is the weakest^34,35^.

To understand how affinity measurements translate to *in vivo* interaction strength, we employed DHFR PCA to quantify the interaction strength of the selected PBDs and peptides. In this assay, the dihydrofolate reductase (DHFR) sequence is split into two fragments, F[1,2] and F[3]. Both fragments are encoded on the same vector in a tail-to-tail orientation, separated by the bidirectional CYC terminator. DHFR F[1,2] and F[3] are under the control of individual CYC promoters and are fused to a flexible linker (F[1,2]-(GGGGS)_4_, F[3]-(GGGGS)_3_-(GGGAS)_1_). The CYC promoter produces a low constitutive expression^36^ of the DHFR fragments, limiting the spurious background signal (Supplementary Fig. 1). The PBD and peptide sequences were codon-optimized for expression in yeast and synthesized. They were cloned individually at the C-terminus end of both DHFR fragments-linker. In a second cloning step, the peptide sequences were cloned in pairs with either F[1,2]-PBD or F[3]-PBD, resulting in two DHFR fragment orientations, i.e. F[1,2]-PBD-F[3]-peptide or F[1,2]-peptide-F[3]-PBD. When both DHFR fragments are colocalized in *S. cerevisiae*, they reconstitute an active enzyme. Under methotrexate (MTX) selection, cell growth is dependent on the number of active DHFR reconstituted, which relies on the interaction strength between the PBD and peptide fused to the fragments (Fig. 1A)^37,38^.

**Figure 1.**
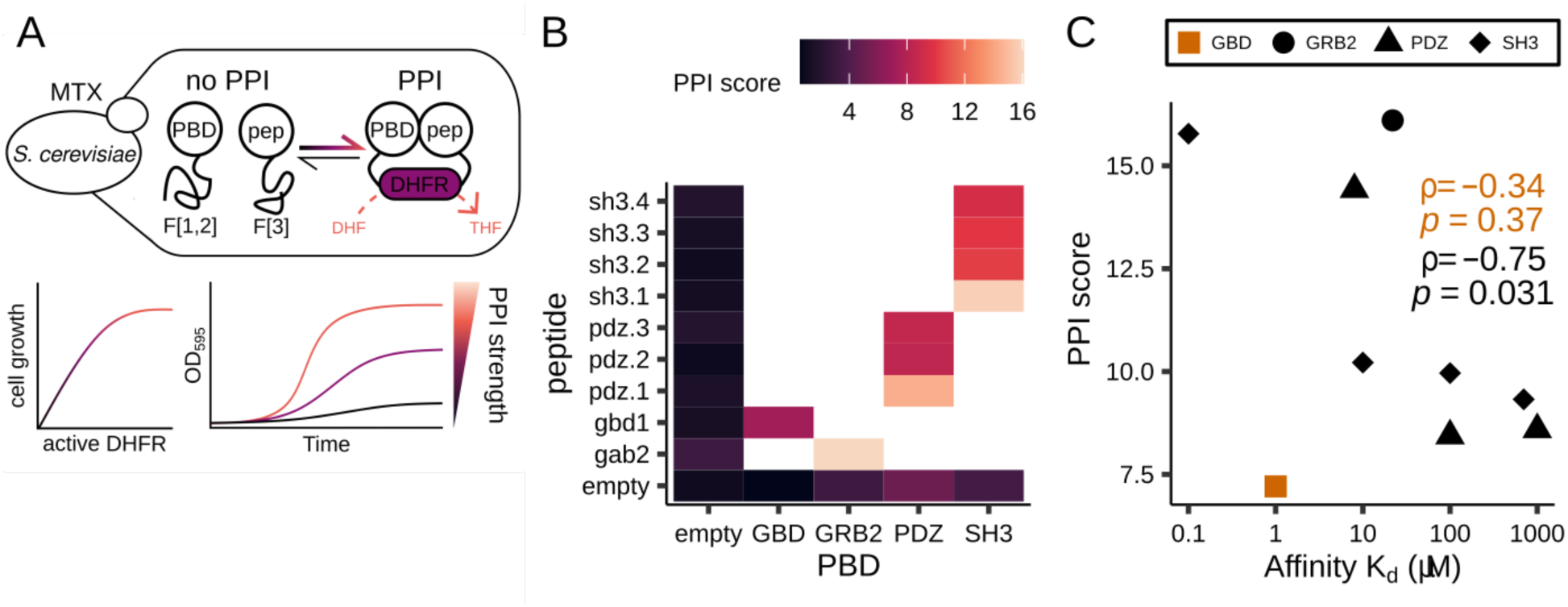
Experimental design and measurement of interactions between PBDs and peptides *in vivo*. A. Schematic representation of the DHFR PCA to measure the interaction strength between a PBD and a peptide in living cells^27^. An interaction between DHFR F[1,2]-PBD and DHFR F[3]-peptide, or vice-versa, allows the DHFR to assemble into its active form. Cell growth is relative to the number of active DHFRs, up to signal saturation. We performed growth measurements under MTX selection and computed the area under the curves to quantify PPI strength. B. Pairs of F[1,2]-PBD and F[3]-peptide tested. The color scale indicates the PPI score of the PBD-peptide pairs. White spaces are untested PBD-peptide pairs. C. Comparison of PPI scores obtained with F[1,2]-PBD and F[3]-peptide orientation with the *in vitro* affinities measured in previous studies^32–35^. The shape of the data points specifies the PBD involved in the binding pair. The Spearman’s rank correlation coefficient and its p-value are labelled when including all data points (orange) or when removing the GBD outlier (black).

To quantify the interaction strength of each binding pair, we assessed the growth of yeast strains transformed with the PBD-peptide containing plasmids under MTX selection (Fig. 1A)^27,32^. We computed the area under the curve (AUC) obtained from the growth measurements of each PBD-peptide combination and calculated a PPI score using the ratio of the growth induced by PBD-peptide pairs over the negative control (empty plasmid, see methods). This relative growth represents the strength of the PCA signal^38^.

The strains expressing either a PBD or a peptide alone show a PPI score very similar to the negative control. As expected, a higher score is measured when a PBD and a peptide are both present (Fig. 1B). Since we cloned every PBD-peptide pair in both DHFR fragment orientations, we assessed the effect of the DHFR fragments fused to each sequence on the PCA signal. We observed that the PCA signal obtained with the peptides fused to F[1,2] is significantly lower than when fused to F[3] for most of the PBD-peptide pairs, with the exception of pairs SH3-sh3.1 and SH3-sh3.3 (Supplementary Fig. 2A). In fact, for most of the sequences tested, the DHFR fragment orientation does not affect the PPI strength rank of the PBD-peptide pairs (Supplementary Fig. 2A). However, we observed an exception for the PDZ-pdz.2 pair. It has a lower signal than the PDZ-pdz.3 pair in the F[1,2]-peptide-F[3]-PBD orientation, which is surprising because the K_d_ of the PDZ-pdz.2 interaction is an order of magnitude lower than the K_d_ of the PDZ-pdz.3 pair (Supplementary Fig. 2A and B). In fact, the DHFR fragment effect is particularly strong on pdz.2 since the F[1,2]-pdz.2-F[3]-PDZ signal is as low as the negative control, but F[1,2]-PDZ-F[3]-pdz.2 signal is similar to F[1,2]-PDZ-F[3]-pdz.3 (Supplementary Fig. 2A). It is possible that the fusion of F[1,2] with pdz.2 leads to a strong intramolecular interaction which prevents the complementation with F[3] and would explain the drastic difference between the two fragment orientations. Since the two DHFR fragments are very different in terms of sequence, pdz.2 might not interact with F[3] in the same way.

Because the F[1,2]-PBD-F[3]-peptide orientation displayed a stronger signal than the opposite orientation, we compared the PPI score obtained by this orientation with affinity values characterized in previous studies^31,33–35^. We detected a wide range of interaction intensity with DHFR PCA, corresponding to signals over multiple orders of magnitude of K_d_ values (Fig. 1C). Although there is a general trend, we did not observe a significant correlation between the PPI scores and the K_d_ values obtained *in vitro* (Spearman’s rank correlation, r = -0.34, p-value= 0.37). The GBD-gbd1 interaction is a visible outlier that causes the lack of correlation (Fig. 1C). Indeed, there is a trend between PPI score and affinity for the SH3 and PDZ domains with their respective binding peptides and the correlation without the GBD-gbd1 becomes significant (Spearman’s rank correlation, r = -0.75, p-value= 0.031). One limitation of this analysis is that the K_d_ values for each PBD were retrieved from different studies and measured in varying ways, which may contribute to this discrepancy. Another factor that could create a disparity between PPI scores and the *in vitro* affinity is the cellular environment, which may affect *in vivo* protein abundance and potentially lead to spurious interactions with yeast endogenous proteins. Other unexpected effects could also diminish the availability of the PBDs and peptides for binding to each other. For these reasons, we performed an assay to quantify the binding availability of each PBDs and peptides *in vivo*.

### PBDs and peptides show different binding availability

Here, we define the binding availability of a protein as the amount of this protein that is able to effectively bind other proteins in the cytosol^30^. This amount can be reduced by the degradation of the protein or by its spurious binding with other partners^25,26^. A strong spurious endogenous binding partner would sequester the heterologous protein and prevent contact with the one we are targeting. In our assays, if a large fraction of a PBD is sequestered by a spurious interaction partner, there is less PBD available to bind with the peptide fused with the complementary DHFR fragment, or vice-versa. This situation could result in a weak PCA signal even if the PBD-peptide interaction characterized *in vitro* is strong. To measure binding availability, we used a binding assay with a non-specific prey, a yellow fluorescent protein (YFP), fused to one of the DHFR fragments^30^. The YFP-DHFR fragment fusions were expressed under a strong promoter inserted in the genome, creating one yeast strain expressing the YFP-F[1,2] and a second strain expressing the YFP-F[3]. We transformed these strains with plasmids expressing either a PBD or a peptide fused to the complementary DHFR fragment (Fig. 2A). The resulting PCA signal reflects non-specific binding events between the YFP and the protein of interest, which is correlated with protein abundance (Fig. 2A)^30^.

**Figure 2.**
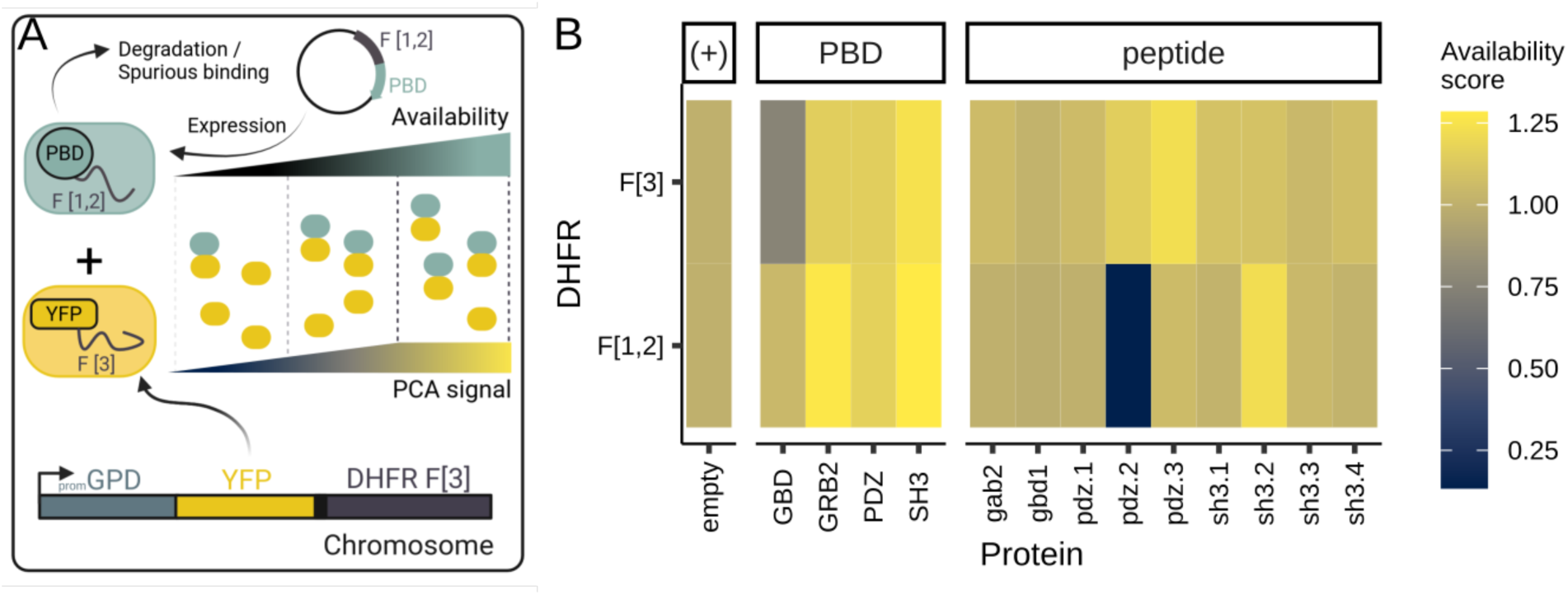
Measurement of binding availability of the PBDs and peptides to be used for scaffolding. A. Schematic representation of the experimental approach to investigate binding availability with DHFR PCA^30^. The PCA signal depends on the non-specific binding between the protein expressed from a plasmid and the YFP encoded in the genome. B. For PBDs or peptides fused to either DHFR fragments, the availability score is shown with the color scale (see methods). The empty plasmid is considered as a positive control (+) in this experiment.

We measured binding availability for all the domains and peptides described previously. We computed the AUC from the growth measurements, and transformed it into an availability score by comparing the AUC values with those from the control expressing the DHFR fragments alone (see methods). We found that most proteins show similar, or higher, binding availability compared to the control (Fig. 2B). One interesting observation is that the pdz.2 peptide showed a similar availability to the control when fused to the F[3], but a very low availability when fused to the F[1,2] (Fig. 2B). The low availability of the F[1,2]-pdz.2 can explain the weak PPI score obtained when tested against F[3]-PDZ (Supplementary. Fig. 2A). We also observed a small but significant difference in the effect of DHFR fragments on the GBD domain, where F[3]-GBD shows less binding availability than F[1,2]-GBD (Student’s t-test, p-value = 1.87x10^-3^, Supplementary Fig. 3). However, the peptide gbd1 fused with either of the DHFR fragments showed similar binding availability to the control. Thus, we were unable to explain the lack of PCA signal in the interaction assay for the F[1,2]-GBD-F[3]-gbd1 pair due to the low binding availability of either the PBD or peptide.

The discrepancy between PPI scores and affinity values is affected by binding availability, but it is insufficient to explain every difference in signal. Following these results, we focused on the SH3 and PDZ domains and their peptides showing similar binding availability to the control and consistent PPI scores with affinity values for the characterization of the fusion of both domains as a scaffold.

### DHFR PCA to detect changes in scaffolding efficiency

Following the characterization of PPI strength and binding availability of PBDs and peptides *in vivo*, our aim was to use this data to better understand their roles in scaffolding efficiency. We constructed scaffolds by fusing the SH3 and PDZ domains for which PPIs and binding availability had been characterized. We created two scaffold architectures by changing the order of the PBDs in the scaffold, namely the SH3-PDZ and PDZ-SH3 orientations. We also tested different linker lengths between the PBDs : direct fusion, a flexible linker (GGGGS)_1_-(GGGAS)_1_ and a longer flexible 2xlinker (GGGGS)_3_-(GGGAS)_1_. We inserted the scaffold variants at the *GAL1* locus. The expression of the scaffold is regulated by a synthetic transcription factor (GEM), which translocates into the nucleus and induces transcription at the *GAL1* promoter when bound to β-estradiol. The synthetic expression system enables the scaffold to be expressed proportionally to the concentration of β-estradiol in the media^28,39^ (Fig. 3A). We validated the inducible expression system by inserting the *mEGFP* coding sequence at the *GAL1* locus and measuring the fluorescence intensity of the cells at different concentrations of β-estradiol (Supplementary Fig. 4).

**Figure 3.**
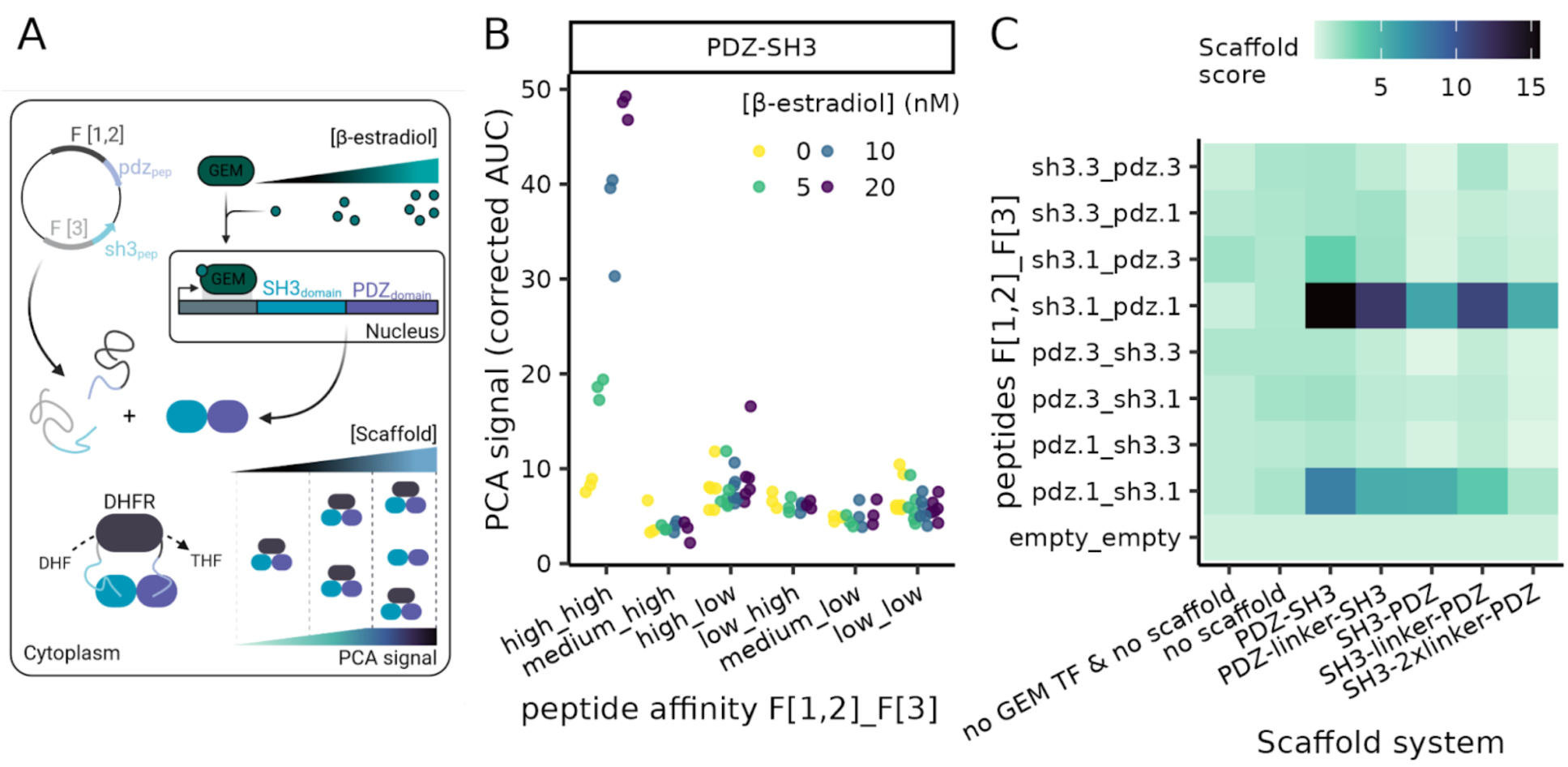
Scaffolding efficiency as measured by DHFR PCA. A. Schematic representation of the experimental system to measure scaffolding efficiency. The scaffold, a fusion of the SH3 and the PDZ domains, is expressed from the genome at an inducible locus controlled by β-estradiol. β-estradiol binds the GEM transcription factor, allowing its translocation in the nucleus and the induction of the scaffold expression^39^. From a plasmid, the DHFR fragments fused to peptides are expressed. PCA signal reflects the ability of the scaffold to colocalize both DHFR fragments by simultaneously binding the peptides. B. For the PDZ-SH3 scaffold architecture, the relation between the PBD-peptide affinity used in the scaffold system (high : K_d_ < 10 μM, medium : 10 μM <= K_d_ < 100 μM, low : 100 μM <= K_d_) and the PCA signal (corrected AUC, see methods) is displayed for different concentrations of β-estradiol. The colors show the β-estradiol concentrations used. C. For different combinations of peptides and scaffold architectures, the scaffold score obtained at 20 nM β-estradiol is shown (see methods).

We reused the same expression vector as in the PPI and binding availability assays, but this time we inserted pairs of peptides binding to the SH3 and PDZ domains in frame with the DHFR fragments. We transformed each plasmid in the strains containing the different scaffold constructs, creating all possible peptide pairs-scaffold combinations. In MTX selection media supplemented with β-estradiol, we expect yeast growth only when the F[1,2]-peptide and F[3]-peptide are simultaneously bound by the scaffold, which allows the complete DHFR to assemble (Fig. 3A). To test how PCA signal depends on the abundance of the scaffold, we can modulate scaffold expression level by changing the β-estradiol concentration, while preserving expression within the dynamic range of the assay. As for the PPI and binding availability assays, we performed growth measurements to quantify the scaffolding efficiency and transformed the AUC into a scaffold score, which is a relative value compared to the negative control.

We found that PBD-peptide affinity, scaffold architecture and scaffold expression level all impact scaffolding efficiency. Expectedly, scaffold scores generally increase as β-estradiol concentrations, and consequently expression level rises, but only when two peptides of high affinity with the PBDs of the scaffold are fused to the DHFR fragments (Fig. 3B, Supplementary Fig. 5). We observe that only the peptide pair with the strongest affinity to the PBDs present in the scaffold (sh3.1-pdz.1) generated a high scaffold score at 20 nM β-estradiol (Fig. 3C). The pairs including at least one lower affinity (K_d_ >= 10 μM) peptide all showed similar signals to the negative control, indicating that scaffolding efficiency is limited by the weakest interaction involved in the system. In addition, we tested combinations of peptides binding to the same PBD (ex. F[1,2]-sh3.1-F[3]-sh3.1). They all showed PCA signals similar to the negative control (Supplementary Fig. 6). This result indicates that two strongly binding and distinct PBD-peptide pairs are necessary for the proper function of the scaffold.

We also found that the scaffold architecture impacts its efficiency. Indeed, the PDZ-SH3 is the most efficient architecture but, when the positions of the PBDs are swapped (SH3-PDZ), scaffolding efficiency is significantly lower for the same peptide pair (F[1,2]-sh3.1-F[3]-pdz.1, Supplementary Fig. 5, Student’s t-test, p-value = 0.031). The length of the linker also unpredictably affects the scaffolding for the F[1,2]-sh3.1-F[3]-pdz.1 pair as reported by DHFR PCA. We observed an intermediate score for the SH3-PDZ architecture, followed by an increase in signal for SH3-linker-PDZ and a lower score for SH3-2xlinker-PDZ. The opposite pattern is observed for the reverse PBD position, i.e. decrease in signal for PDZ-linker-SH3 compared to PDZ-SH3 (Fig. 3C).

In order to ensure that different architectures led to a similar protein abundance, we examined protein abundance by western blotting. We added a FLAG epitope to each scaffold, at either their N- or C-terminus, and measured their abundance at an induction level of 20 nM β-estradiol (Supplementary Fig. 7). We observed similar abundance levels for all scaffold architectures with the C-terminal FLAG (Supplementary Fig. 7B). However, we detected a slight degradation of the scaffolds with a PDZ-SH3 orientation compared to the SH3-PDZ orientation, but it is not sufficient to reduce the scaffolding efficiency since PDZ-SH3 is the most efficient scaffold architecture. When tagged at their N-terminal, the scaffolds with a SH3-PDZ orientation showed variation in their abundance levels. The SH3-PDZ scaffold has a lower abundance compared to SH3-link-PDZ scaffold, which could explain its lower efficiency (Supplementary Fig. 7C). Overall, the design features tested impact scaffolding efficiency and even the scaffold abundance level, which highlights the relevance to characterizing in depth variants of a scaffold before choosing one for practical applications.

### Characterizing new PBD-peptide candidates *in vivo*

Our PPI and scaffolding PCA experiments showed that a minimal PPI strength is needed for the PBD-peptide pairs to build an efficient scaffold in yeast. Thus, we explored new candidates of PBD-peptide pairs that could show similar PPI intensity to PBDs already used in scaffolds. To select domains with distinct binding specificity, we searched the Peptide Recognition Module Database (PRM DB)^40^. The three main PBD families present in the PRM DB are the SH3, the PDZ and the WW repeating motif (WW) domains. These PBD families are known to have a wide range of affinities to their binding peptides and diverse specificities^18,41^. As described above, PBDs previously used in scaffolds were an SH3 and a PDZ domain; thus, we can be confident that these PBD families can create strong enough PPI to be of interest for synthetic scaffold applications.

In the design of synthetic systems, it is important to use PBDs that have orthogonal specificity in order to avoid crosstalk between synthetic circuits. Thus, we used the position weight matrices (PWMs) compiled in the PRM DB to select PBDs with diverse specificities. We computed the minimal average absolute difference (AAD) in amino acid frequency between the PWMs as a metric to compare specificity within a PBD family (Figure 4A, 4B, see methods)^42^. An AAD of 0 indicates a perfect agreement between the PWMs and an AAD of 0.1 indicates a strong divergence between the PWMs. By computing the minimal AAD between all PWMs of a PBD family, we identified clusters of specificity as groups of PBDs which would bind similar peptides (Supplementary Fig. 8). To select orthogonal PBD-peptide pairs, we chose one PBD per cluster. We validated that their PWMs report divergent specificity (Supplementary Fig. 9). We selected seven SH3 (PBD 131, 152, 214, 246, 250, 299, 304), three PDZ (PBD 363, 366, 385) and two WW (PBD 25, 61) domains for characterization *in vivo*. Apart from the SH3 domain of Sho1 (PBD 152), all of these PBDs are exogenous to *S. cerevisiae* and, to our knowledge, their binding has not been characterized in this host. For peptide selection, we based our design again on the PRM DB by using the most abundant peptides in the phage-display data for each PBD. In addition, we extracted from each PWM the peptide sequence encoding the top-scoring amino acid at each position. These criteria allowed us to identify 53 PBD-peptide pairs for individual characterization in yeast.

**Figure 4.**
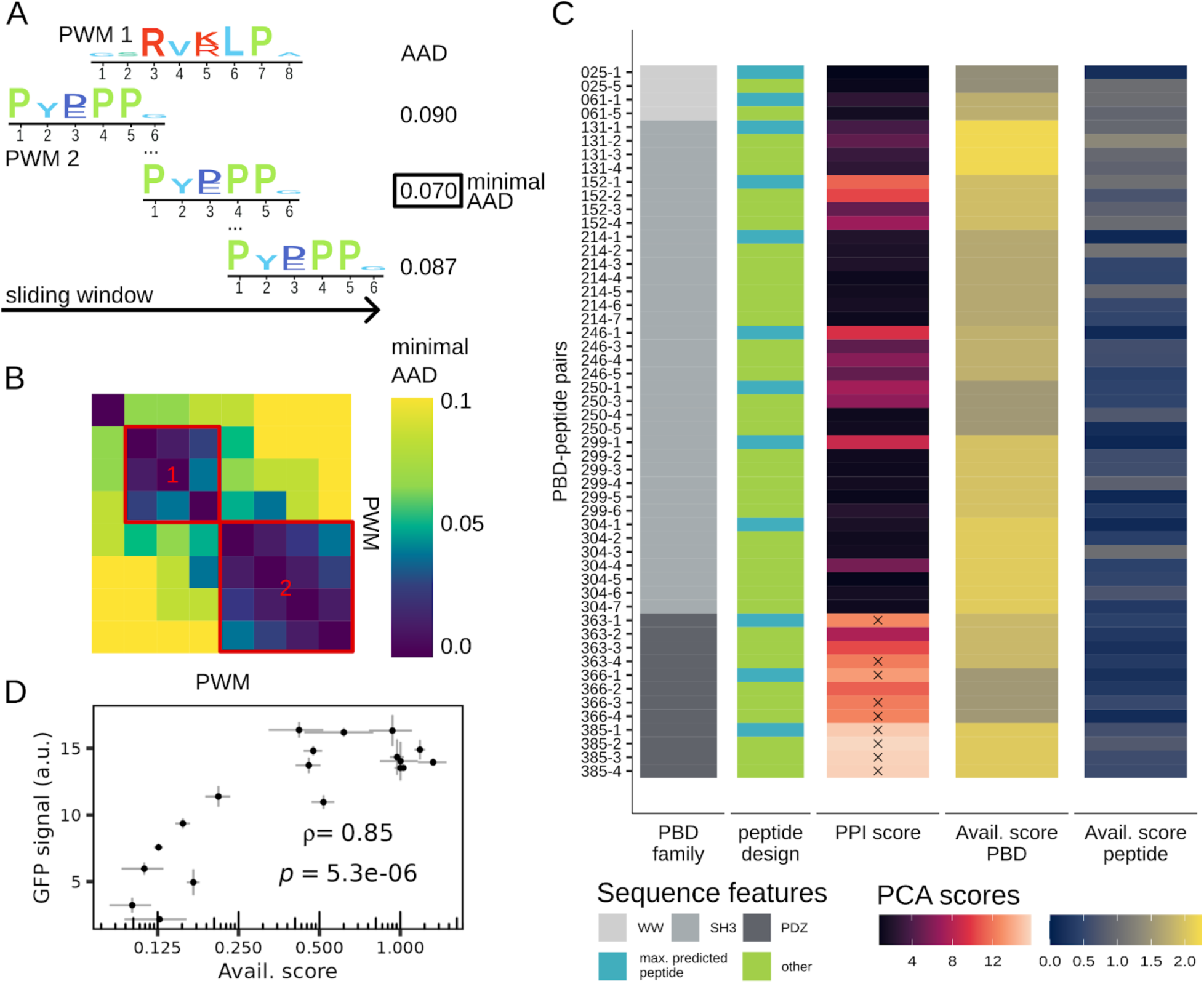
Selection of novel PBD-peptide pairs and characterization of their PPI strength and binding availability. A. Schematic representation of the method used to select the PBDs using the PWMs from the PRM DB^40^. We computed the AAD between two PWMs following a sliding window approach where at least half the positions of one PWMs were considered. We kept the minimal AAD value for each comparison. B. The minimal AAD between the PWMs was used to identify clusters of specificity in each PBD family. The identification of specificity clusters allowed us to select PBDs with divergent specificity (Supplementary Fig. 8) C. PPI strength and binding availability PCA scores for the selected PBDs-peptide pairs associated with their sequence features. The X identifies the PBD-peptide pairs that have stronger or similar PPI scores (> 12) than the pairs used in previously characterized scaffolds. D. The comparison between availability scores and fluorescent intensity detected when the mEGFP is fused to a peptide (arbitrary units). The Pearson correlation coefficient and its p-value are labelled on the plot. The standard deviation between the replicates (n = 3) is shown by the error bars for each data point.

We codon-optimized the PBD and peptide sequences for expression in *S. cerevisiae* and ordered them as synthetic DNA. We then inserted the PBD sequences in frame with the F[1,2]-linker and the peptide sequences in frame with F[3]-linker, paired or alone, in the same vector as for the previous assays. When the PBD and peptide are paired, we used the DHFR fragment orientation, which gave the strongest PPI signal in our previous characterization i.e. F[1,2]-PBD-F[3]-peptide. We used the DHFR PCA approach with growth curve analysis for the characterization of the PPI strength and binding availability of the candidate PBD and peptide sequences^27,30^.

We observed that not all PBD-peptide pairs showed an interaction *in vivo* (Figure 4C). In fact, only eight PBDs from the 12 selected produced detectable PPI with DHFR PCA, which do not include any of the WW domains. We detected interactions for the SH3 domain of Sho1 (PBD 152) with its designed peptides, which indicates that DHFR PCA is able to detect biologically relevant PPIs since this is an endogenous PBD. When a PBD produces a PCA signal, its strongest PPI is nearly always observed with the best binding peptide predicted by the PWM (7/8 PBDs, Figure 4C). This observation indicates that the specificity information obtained by phage display is transposable to synthetic interaction *in vivo*. Since the phage-display assay does not capture interaction strength, it is not surprising to observe an absence of PCA signal for some PBDs (PBD 25, 61, 214, 304). Our experiment reveals that the binding properties of proteins should be investigated in a cell host, even if extensive *in vitro* characterization was performed since the portability of PPI *in vivo* can be unpredictable.

Out of the 12 PBDs characterized, three PBDs showed a PPI strength signal similar to the PBD-peptide pairs enabling a strong scaffolding signal (PDZ-pdz.1 and SH3-sh3.1), with at least one peptide. In the PCA experiment with the peptides colocalizing to the scaffold, we observed that the PCA signal dropped when one PBD-peptide pair had a PPI score inferior to 12. Thus, we use this value as a threshold to identify the PBD-peptide pairs that are of interest for synthetic scaffold design. Namely, the PBDs 363, 366 and 385 showed PPI signal with a peptide above this threshold (Figure 4C).

When comparing the PPI scores and the binding availability scores, we do not observe an association between the two assays (Figure 4C). All PBDs showed a higher binding availability than the control, which indicates that the absence of PPI for some PBDs is not related to low availability of these domains. However, most peptides show lower binding availability than the control, which could be explained by enhanced degradation of the F[3]-peptide fusion protein or that they are sequestered by some spurious binder. Thus, we investigated the effect on protein abundance of the peptides by fusing some of them to the mEGFP-linker(GGGGS)_3_-(GGGAS)_1_ regulated by the CYC promoter and terminator on a plasmid. We measured the fluorescent intensity of cells transformed with plasmids expressing the mEGFP-peptide variants using flow cytometry. We observed a significant correlation between the mEGFP-peptide abundance and the F[3]-peptide availability scores obtained by DHFR PCA (Figure 4D, Pearson correlation coefficient = 0.85, p-value = 5.3x10^-6^). This observation indicates that some peptides reduce the abundance of the protein they are fused to. However, even peptides with a low binding availability can be involved in the strongest interactions. For example, the strongest binder to PBD 246, peptide 246-1 (PPI score = 9.38), has the lowest binding availability (peptide Avail. score = 0.13) detected in the 246 associated peptides and displayed a low fluorescence signal when fused to the mEGFP (mean GFP signal = 2.18). This observation indicates that even at lower abundance, a peptide can interact with its designed PBD and generate a PCA signal. If the peptide fusion does not lead to reduced protein abundance, it is possible that the low availability peptides bind spuriously to an endogenous protein or that they create an intramolecular interaction with the DHFR fragment, thus interfering with the DHFR complementation and the binding availability PCA signal. Then, when the PBD is co-expressed with the peptide, the stronger interaction between the PBD-peptide pair could displace the spurious interaction, which would induce the PCA signal. Even if further characterization of the peptides is needed to understand the mechanism behind the variability of binding availability scores, we were able to identify sequences that would be suitable to build efficient scaffolds. Indeed, we have found three exogenous PBDs (363, 366, 385) which, when combined with peptides, produced PPI scores above the threshold identified (PPI score > 12), making these domains good candidates for scaffold design.

## Discussion

Most past experiments measuring scaffolding efficiency have relied on pathway output to quantify a scaffold’s ability to induce colocalization of the targeted proteins, which has limited the throughput for testing various scaffold architectures and sequences^24^. Thus, there was a need to develop a strategy to study scaffolding efficiency directly with an easy-to-read output. Using DHFR PCA, we characterized the PPI strength and binding availability of PBD-peptide pairs. We showed that the position of the PBDs, linker length, PBD-peptide affinity, and scaffold expression level play a significant role in the scaffolding efficiency. Additionally, we characterized PBD-peptide pairs that were not used in scaffolds previously.

A growing number of synthetic protein scaffolds are available due to their high potential for pathway optimization. PBD-based scaffolds are only one of the designs available. Other protein scaffold designs use relatively large chimeric proteins, such as the cohesin-dockerin systems (∼70 a.a.)^43–45^, that may be damaging for enzyme activity^1,46^. A limitation of the cohesin-dockerin systems is their calcium dependency^47^, which makes them less efficient for intracellular assembly because cytosolic calcium concentration is low in yeast^48^. Novel designs use inducible protein condensates, which do not yet offer control on interaction specificity between the proteins involved in the condensate formation^49^. Thus, the PBD-peptide scaffold strategy is an interesting option since a small peptide (∼10 a.a.) fusion on an enzyme of interest is expected to have limited damaging effects on its activity and folding. Moreover, the intrinsic specificity of the PBDs should ensure specific recruitment in the cytosol. However, as we have seen with F[1,2]-pdz.2, peptides can affect protein availability and impact interaction strength. We also observed that the protein fused to a PBD can change its binding availability. For instance, we saw that the GBD domain’s availability can change depending on the attached DHFR fragment and that its interaction strength with the gbd1 peptide in yeast is lower than expected based on its K_d_. These observations could explain why the synthetic scaffold using the pair GBD-gbd1 showed limited yield increase in yeast^23^ compared to what has been shown in bacteria.

One approach to increase synthetic pathway efficiency that is often compared with the use of synthetic scaffold is the direct fusion of proteins of a pathway. At times, directly fusing enzymes showed increases in product yield in comparison to having the enzymes separated and, in some cases, to using a scaffold^21,50,51^. However, directly fusing the enzymes can sometimes be unfavorable for protein folding or enzymatic activity. The PBD scaffold approach was shown to be more efficient than enzyme fusions, for instance, in the biosynthesis of resveratrol^23,52^. Indeed, depending on the pathway of interest, a scaffold could be more efficient than a fusion, or vice versa.

Additionally, the effect of the ratio of enzymes involved in a pathway is a main determinant of pathway efficiency, since the enzymes in the pathway may have different kinetic parameters and could cause flux imbalance leading to accumulation of intermediates. If a toxic intermediate accumulates, it may result in the diffusion of the intermediate in the cytoplasm and cause damage to the host. With a scaffold strategy, the enzyme ratio is not limited by sequence fusion to 1:1^53^. Indeed, we can adapt the number of each PBD in a scaffold to optimize the colocalized enzyme ratio and to address the flux imbalance by recruiting the flux-limiting enzyme in higher amounts, locally increasing the rate of its reaction^17,23^. Then, we can change PBDs contributing to the scaffold to improve compatibility with the pathway of interest. For instance, a specific peptide fusion could compromise the enzymatic function or protein availability of one enzyme, while a change of peptide sequence could resolve the issue. Since very few PBD-peptide pairs have been characterized in a scaffold so far, optimizing a scaffold by changing the PBD and peptide sequences is hard. With the DHFR PCA, we resolved this bottleneck by characterizing other pairs of PBD-peptide *in vivo*, providing alternative PBD-peptide sequences. The PPI assay showed that there can be discrepancies between *in vitro* and *in vivo* binding strength, and the peptide abundance measurements showed that some peptides lead to the degradation of the protein they are fused to. These two observations underscore the importance of characterizing heterologous sequences *in vivo* before utilizing them as tools for synthetic biology applications. Once the toolbox for scaffold design is expanded to more PBD-peptide pairs, the scaffold strategy might outperform the fusion design for most pathways by fully taking advantage of its versatility.

Some limitations of our approach to quantify scaffolding efficiency are connected directly with the number of DHFR fragments available and with the expression system we used. As shown by Dueber et al. 2009^17^, we found that scaffold expression levels can be used to optimize the output signal induced by a scaffold. To minimize background signal, we used a constitutive low-expression promoter to express the PBD and peptides fused to the DHFR fragments (Supplementary Fig. 1). A low-expression PCA system is favorable because it ensures detection below signal saturation and it has a limited metabolic burden. If the expression level of the DHFR fragment-peptides fusion had been higher, we might have detected scaffolding signals for weaker binding PBD-peptide pairs. In the future, higher expression of the scaffold and DHFR fragment-peptides fusion could be tested to evaluate the lower limit of detection of the assay and the effect of metabolic burden on cell growth. Another potential limitation to our assays is that the DHFR PCA used to characterize scaffolding efficiency is constrained to two-component scaffolds, which is restrictive because most synthetic pathways involve more than two enzymes. However, our approach can be used to test if two PBDs are compatible for scaffold design, and then various pairs could be combined. Indeed, the folding of some PBD fusions could hinder one binding site of the scaffold and lead to an inefficient scaffold even if the PBD-peptide pairs showed strong interactions. Also, the effect of linker properties^54^ between the PBDs could be further investigated using our method, which has not been done, to our knowledge, for PBD-based scaffolds.

To further develop the PBD scaffold strategy, the DHFR PCA for scaffold characterization can be adapted for a competition experiment using libraries of PBD-peptide pairs and scaffolds. We could detect the best combinations by following the frequency of all sequence variants in MTX selection with deep sequencing^19^. A competition experiment may help in identifying more PBD-peptide binding pairs suitable for scaffold design and more efficient scaffold architectures with the aim of expanding the toolbox for synthetic scaffold design in yeast. Although many parameters of synthetic pathway design may come into play for pathway efficiency, the use of an efficient synthetic scaffold has the potential to be an important building block for effective bioprocesses.

## Author contributions

P.L., A.K.D. and C.R.L. were involved in the experimental design. P.L. and A.K.D. designed molecular biology material for this study. P.L. performed the experiments. P.L. completed the analysis and wrote the first draft of the manuscript. P.L., A.K.D. and C.R.L. contributed to writing, revising and approving the final version of the manuscript. C.R.L. and P.L. acquired the funds to support this project. C.R.L. supervised this work.

## Acknowledgments

We would like to thank Guillaume Diss who provided key materials and comments for the experimental design. We are thankful for the comments of Soham Dibyachintan, David F. Jordan, and Isabelle Gagnon-Arsenault on the manuscript, and for the support of all other members of the Landry Lab. Some figures were produced using BioRender.

## Funding

This work was funded by a Canadian Institutes of Health Research Foundation grant number 387697 and 525827. C.R.L. holds the Canada Research Chair in Cellular Systems and Synthetic Biology. P.L. was supported by an Alexander Graham Bell PhD scholarship from NSERC (BESC D) and a Leadership and Sustainable Development Scholarship from Université Laval.

## Data availability

All growth measurements, flow cytometry data, original western blot images, and supplementary material including the code for data analysis and figure generation is available on the Github repository (https://github.com/Landrylab/Lemieux_et_al2025). The strains and plasmids are available upon request. Supplementary File 1 contains the supplementary methods and Supplementary File 2 contains the Supplementary Tables 1 to 6.

## Conflict of interest

The authors declare no conflict of interest.

## Material & Methods

The following is a summary of the methods and more details, including media composition and detailed PCR cycles, can be found in the supplementary methods (Supplementary File 1).

### Plasmid construction

The destination plasmid (pGD110), kindly donated by Guillaume Diss^32^, allows for the expression of the murine dihydrofolate reductase fragments, DHFR F[1,2] and DHFR F[3], encoded on opposite DNA strands in a tail-to-tail position. Both fragments are under the control of individual CYC promoters and one bidirectional CYC terminator. The DHFR fragments are fused to flexible linkers in C-terminal (DHFR F[1,2]-(GGGGS)_4_, DHFR F[3]-(GGGGS)_3_ -(GGGAS)_1_). pGD110 was digested with BamHI-HF or HindIII-HF (NEB) for the insertion of sequences in frame with, respectively, the F[1,2] or F[3], in N-terminal. The PBDs, gbd1 and gab2 DNA sequences were codon optimized for expression in *S. cerevisiae* and synthesized (Integrated DNA Technology, IDT, Supplementary Table 1). They were amplified with 40-pb homology arms to pGD110 for both DHFR fragment fusion using the standard PCR reaction mix and cycle (See detailed methods, Supplementary File 1) and purified using magnetic beads (Axygen AxyPrep Mag PCR Clean-up SPRI beads). Gibson assembly was performed using 50 ng of digested plasmids with an insert molar ratio of 1:3 (backbone : insert) in a 10 μL reaction volume^55^. The reactions were incubated at 50°C for 60 minutes and 5 μL of the reactions were transformed in MC1061 chemo-competent cells. All transformations were plated on solid 2YT medium with ampicillin (100 μg/ml, see all media composition in Supplementary File 1). The other peptide coding sequences were codon optimized for expression in *S. cerevisiae* and synthesized as oligonucleotides (Eurofins). They were inserted downstream of both DHFR fragments paired with a PBD or alone, by site-directed mutagenesis using the standard PCR mix and cycle. The mutagenesis products were digested with DpnI and 5 μL were used for transformation in MC1061 chemo-competent cells. All the synthetic DNA used for the plasmid construction is described in the Supplementary Tables 1 and 2. All plasmids were validated by Sanger sequencing (CHUL sequencing platform) and they are listed in Supplementary Table 3. The PBD and peptide sequences are listed in Supplementary Table 4.

### Strain construction

The strains used for the binding availability assay express a YFP-linker(GGGGS)_2_-DHFR fragment under the GPD promoter control. First, the pGPD-YFP-linker(GGGGS)_2_-DHFR fragment, F[1,2] or F[3], sequence was assembled on a plasmid. Each fragment was PCR amplified with 40-pb homology to each neighbor fragment. The GPD promoter sequence was amplified from pAG416-GPD-ccdb^56^, the linker-DHFR fragments were amplified from pAG25-DHFR F[1,2]-NATMX and pAG32-DHFR F[3]-HPHMX^27^, the YFP sequence was amplified from 34a_mVenusNB (Michael Springer Lab). The backbone was amplified from pKB1. All fragments were amplified using the standard PCR mix and cycle. Gibson assembly was performed as described above. The resulting plasmids, pPL1 (pKB1-YFP-linker(GGGGS)_2_-DHFR F[1,2]-NATMX) and pPL2 (pKB1-YFP-linker(GGGGS)_2_- DHFR F[3]-HPHMX), were validated by Sanger sequencing. pPL1 and pPL2 were transformed using standard lithium acetate yeast transformation^57^ in BY4741 (*his3 leu2 ura3 met15*) competent cells and plated on selective media (YPD+NAT or YPD+HYG) creating the strains PL0017 and PL0019, respectively. The strains were then transformed with a plasmid expressing a PBD or a peptide fused to either of F[1,2] or F[3]. The pGD110 is also transformed in both strains to assay the basal spurious binding between the F[1,2] and F[3] expressed from the plasmid and the YFP-DHFR fragments. The comparison between the PCA signal obtained from growth measurements of pGD110 and the plasmids expressing a PBD or a peptide reports the relative binding availability of the added sequence to the plasmid.

To create the strains to perform the scaffold assays, we retrieved a strain expressing the GEM transcription factor from a previous study (PL0001)^28,39^. The scaffold sequences were synthesized (IDT) and PCR amplified with 40-pb homology arms to the *GAL1* locus in 5’-terminal and to the NATMX cassette in 3’-terminal using PCR4. The resulting DNA fragments were integrated in strain PL0001 by homologous recombination using standard lithium acetate yeast transformation^57^ and plated on selective media (YPD + NAT). The scaffold strains were validated by PCR and Sanger sequencing (See detailed methods, Supplementary File 1). All the synthetic DNA and plasmids used for the strains construction are described in Supplementary Tables 1-3 and the constructed strains are listed in Supplementary Table 5.

### DHFR PCA assay and analysis

The DHFR PCA protocol is based on Tarassov et al. 2008^27^. Yeast strains transformed with a plasmid were grown in liquid synthetic media (SC-ura) then diluted to an optical density (OD) at 595 nm of 0.1 in DHFR PCA selection media and incubated in transparent polystyrene 96-well plates (Greiner Bio-One). When needed, β-estradiol (Sigma-Aldrich) diluted in ethanol (GreenField Global) was added to the DHFR PCA selection media at concentrations ranging from 0 nM to 20 nM. Cell growth was measured in three or four replicates without shaking by measuring the OD at 595 nm in 15-minute intervals over two days of incubation at 30°C in a BioTek Epoch 2 plate reader (Agilent). The median of the first 5 measurements was subtracted from all the following measurements to correct for variation in initial cell inoculate density. Empirical area under the curve (corrected AUC) was computed using the first 40 hours of growth. The corrected AUC metric was chosen over the derivative growth rate because the corrected AUC showed weaker background signal. The PCA scores (PPI, availability and scaffold scores) were obtained by calculating the ratio of the sample average corrected AUC on the control average corrected AUC (F[1,2]-empty-F[3]-empty) in the same strain background and growth condition.

### PBD and peptide sequence selection from the PRM DB

The Peptide Recognition Module database (PRM DB)^40^ was used for the choice of peptide binding domains (PBD) and their paired selection of peptides. The position weight matrices (PWMs) of the PRM DB were filtered by the number of peptides used to generate them (# peptide > 30). We compared the remaining PWMs of the same PBD family, i.e. SH3, PDZ and WW domains, with each other by computing the average absolute difference (AAD) between each PWMs. The AAD is the sum of the absolute frequency deviations for all amino acids, divided by 20 where 0.0 is a perfect match between PWMs and 0.1 is the maximal divergence^42^. Since the PWMs we compared were not necessarily of the same length, we computed the AAD for every possible comparison between two PWMs where half of the positions of the shorter matrix were compared to the longer one in a sliding window manner. We kept the minimal value of AAD as a value of divergence. Minimal AAD allowed us to calculate clusters of specificity present in the PRM DB (Supplementary Figure 8) and to select PBD sequences which have divergent specificity (Supplementary Figure 9). We selected two WW, three PDZ and seven SH3 domains for protein-peptide interaction *in vivo* assay. From the PRM DB, we selected the most abundant peptides obtained by the phage-display assay (named NGS) and their Elisa validated peptides (named HAL) for PPI validation. We also generated the maximal scored peptide according to the PWM for each PBD by selecting the top scored amino acid at each position (PWM_top). All PBD and peptide sequences were codon optimized for expression in *S. cerevisiae*. The PBD sequences were synthesized in pairs on a vector (Table S1, IDT) and the peptide coding sequences were synthesized as oligonucleotides (Table S2, IDT). The PBD sequences were cloned in frame with the F[1,2]-linker in pGD110 using the same approach as described in the plasmid construction method. The oligonucleotides encoding the peptide sequences were used as primer forward in a PCR (PCR5, see Supplementary File 1 - Detailed methods) on pGD110 to create the gene fragment to insert in frame of F[3]-linker of pGD110. The resulting DNA fragments were purified with magnetic beads and used as inserts in a Gibson assembly, followed by a transformation in MC1061 chemo-competent cells (as previously described). Most plasmids (119/124 pGD110 variants) were validated by colony PCR and Sanger sequencing (CHUL sequencing platform). All plasmids are listed in Supplementary Table 3 and the peptide sequences and are listed in Supplementary Table 4.

### GFP-peptide abundance measurements by flow cytometry

The pPL9-variants, pGD110 backbone modified to encode the mEGFP-linker-peptide (see Supplementary File 1 - Detailed methods), were transformed in the strain BY4741 and plated on selective media (SC-ura). The transformed strains were grown overnight in triplicates in SC-URA. Cultures were diluted in fresh media (OD ∼0.15) and grown to exponential phase (OD = 0.5 - 0.7), then they were diluted to 0.05 OD in sterile water for cytometry measurements. The fluorescent intensity of 5000 cells per replicates was measured using a Guava EasyCyte HT cytometer (blue laser, λ = 488 nm). The median fluorescent intensity of each replicate was computed and used to compare the pPL9-variants

## Abbreviations

DHFR: Dihydrofolate reductase
GBD: GTPase protein binding domain
PBD: Peptide binding domain
PCA: Protein-fragment complementation assay
PDZ: PSD95, Dgl1, Zo-1
PPI: PBD-peptide interaction
SH3: Src Homology 3

---

**Supplementary Figure 1.**
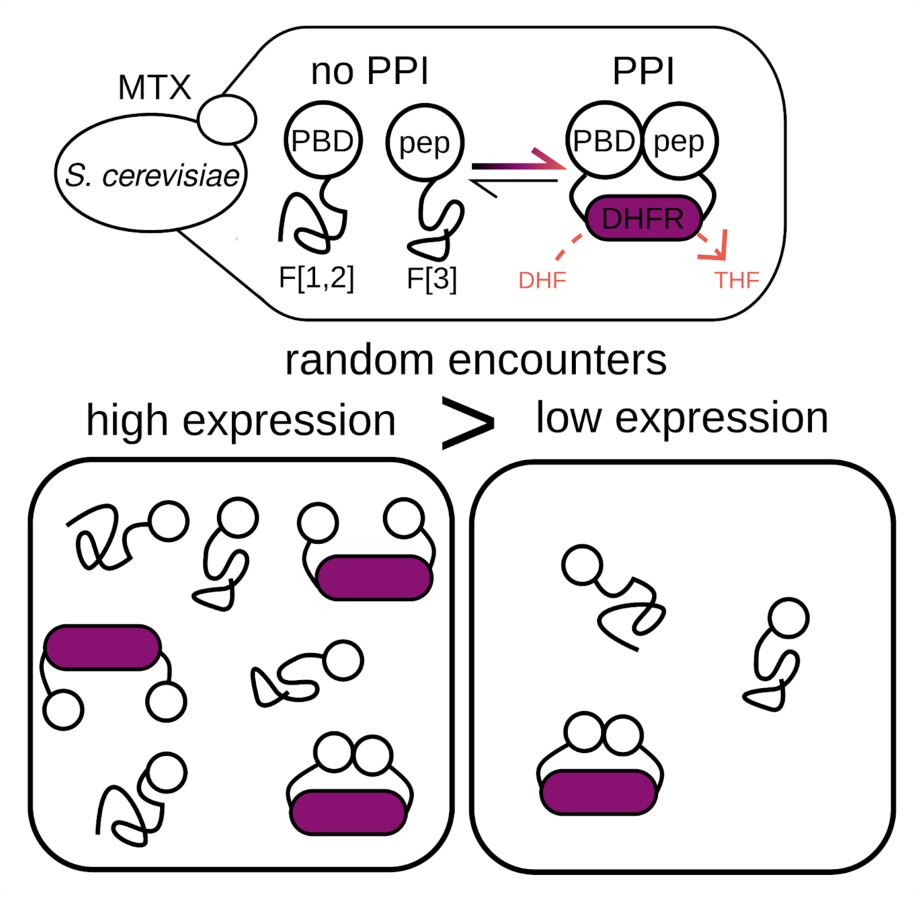
Effect of the expression level of the DHFR fragments on PCA background signal. Schematic representation of the DHFR PCA to measure the interaction strength between a PBD and a peptide. The random encounters between the proteins fused to the DHFR fragments can form an active DHFR, generating THF and thus growth, generating a PCA signal. If the expression level of the DHFR fragments is high, more random encounters happen, thus increasing the PCA signal even if the interaction between the two proteins fused is weak. Also, DHFR PCA can reach signal saturation faster when the background signal is high, reducing the dynamic range of the assay. With a lower level of expression of the DHFR fragments, there is less random encounter and the signal range is reflecting the interaction strength between the proteins fused.

**Supplementary Figure 2.**
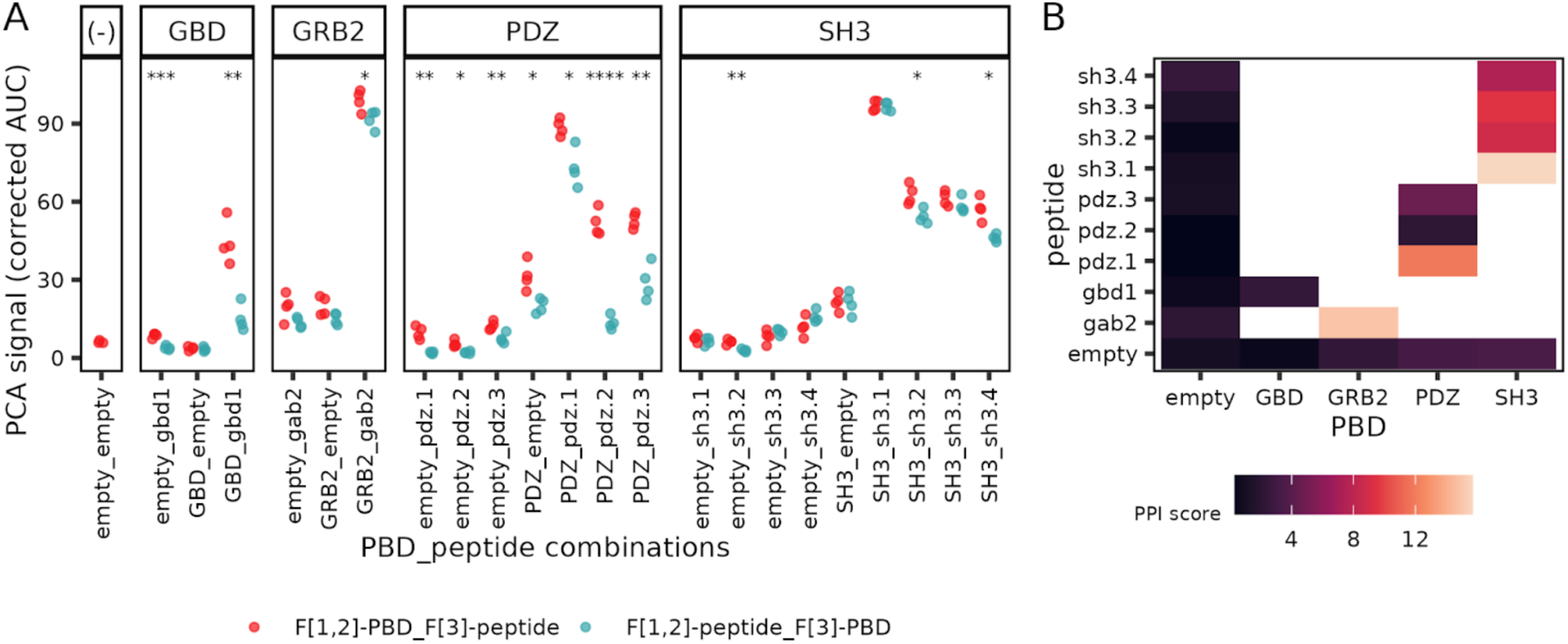
Detailed results of the DHFR PCA PPI assay. A. For each PBD and peptide, paired or alone, the corrected area under the curve of the DHFR PCA assay is shown for each replicate. Empty refers to no sequence inserted as the PBD or the peptide. The color represents the DHFR fragment orientation of the combination. Student’s t-tests were performed between the two orientations for each sequence combination and the p-values significance levels are labeled above each comparison ( * : p <= 0.05, ** : p <= 0.01, *** : p <= 0.001, **** : p <= 0.0001). B. Pairs of F[3]-PBD and F[1,2]-peptide tested with DHFR PCA. The color intensity indicates the interaction strength between the PBD-peptide pairs compared to the negative control (PPI score, see methods).

**Supplementary Figure 3.**
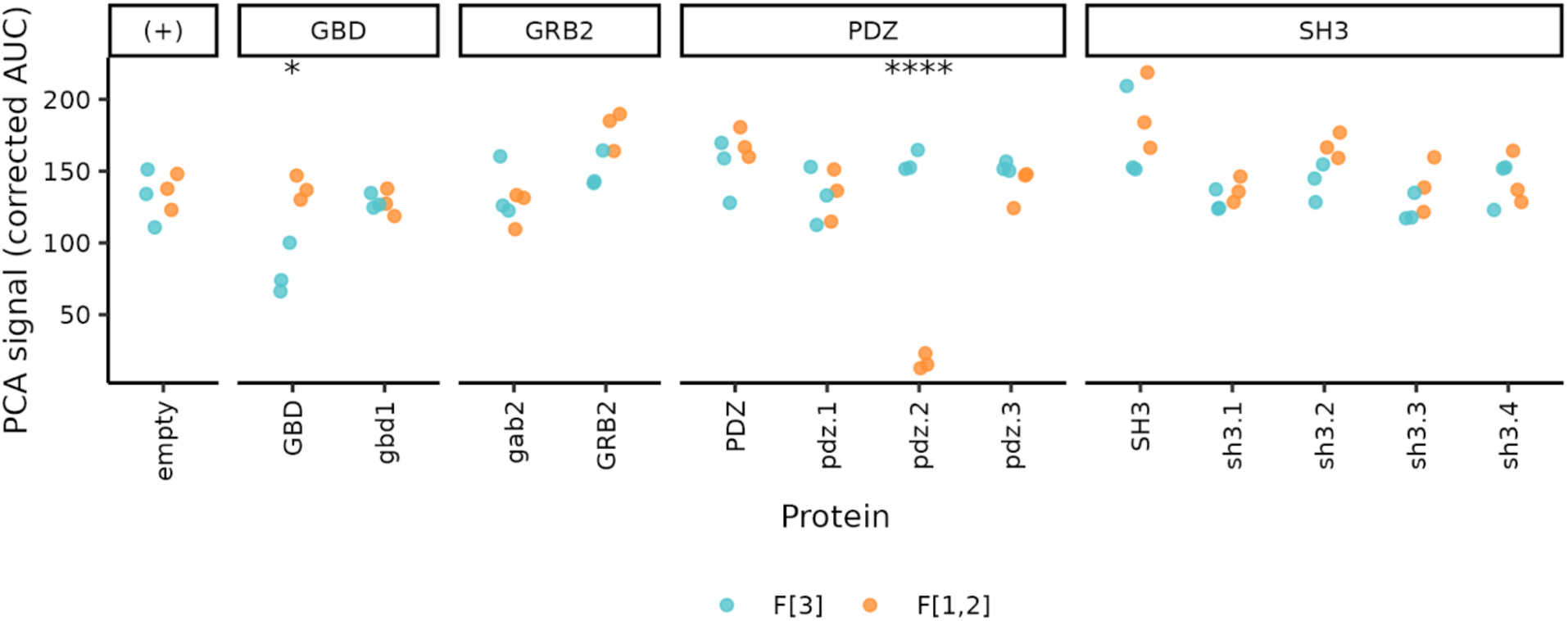
Detailed DHFR PCA results of the binding availability assay. The corrected area under the curve of the availability assay is shown for each sequence tested in triplicate. The color shows the DHFR fragment fused to the proteins. Student’s t-tests were performed to assay the DHFR fragment effect for each sequence and the p-values significance levels are labeled above each comparison ( * : p <= 0.05, **** : p <= 0.0001). The empty plasmids are considered as positive controls (+) in this experiment.

**Supplementary Figure 4.**
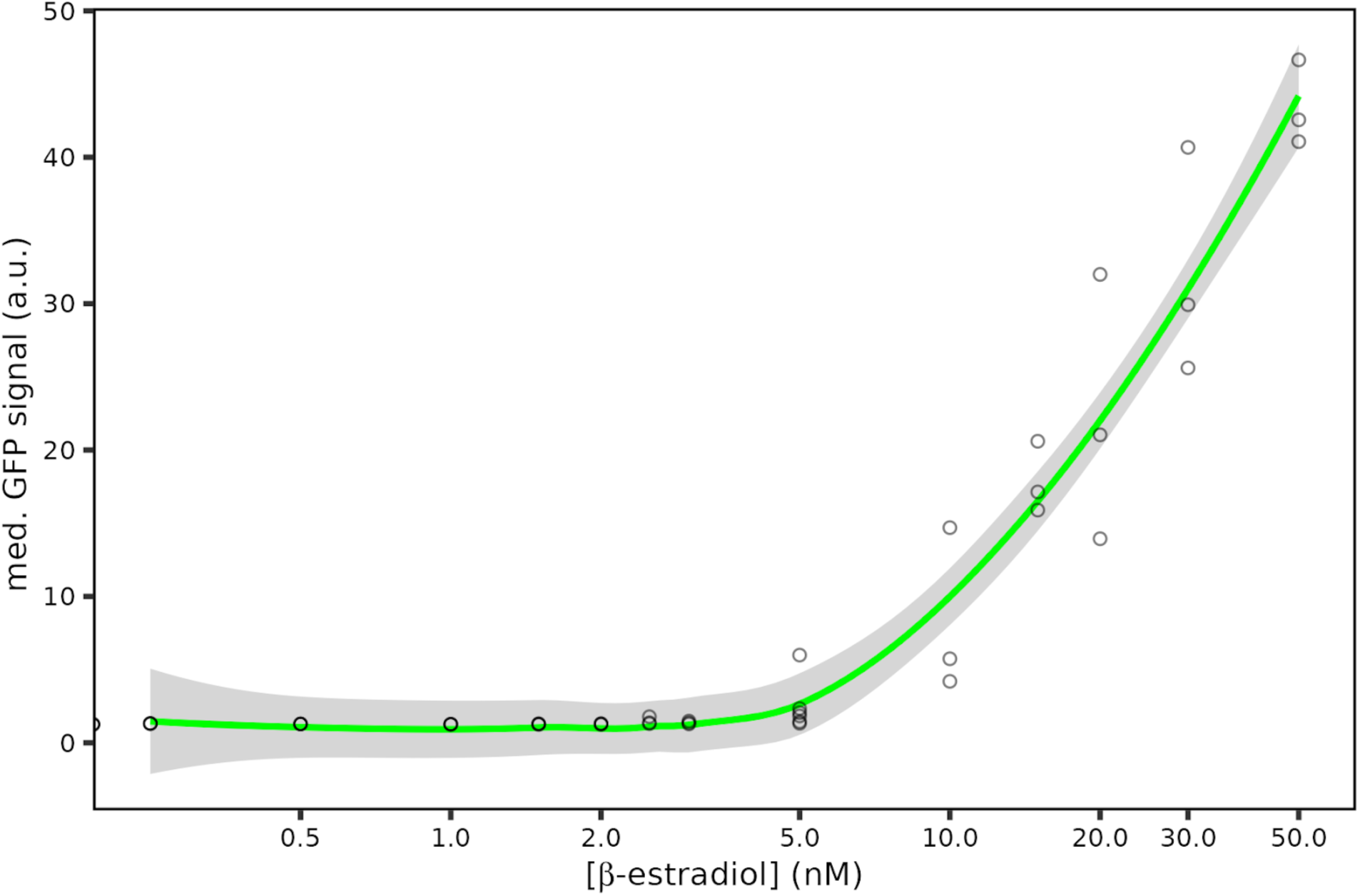
Validation of β-estradiol induction system. The mEGFP coding sequence was inserted at the locus under the control of β-estradiol induction, *GAL1*, and we measured the fluorescent intensity of the cells in exponential growth phase (n = 5000) at β-estradiol concentrations ranging from 0 to 50nM in at least 3 replicates. Each data point shows the median fluorescent intensity of a replicate. The green line represents the local polynomial regression of the median GFP signal over β-estradiol concentration and the shaded area is its 95 % confidence interval.

**Supplementary Figure 5.**
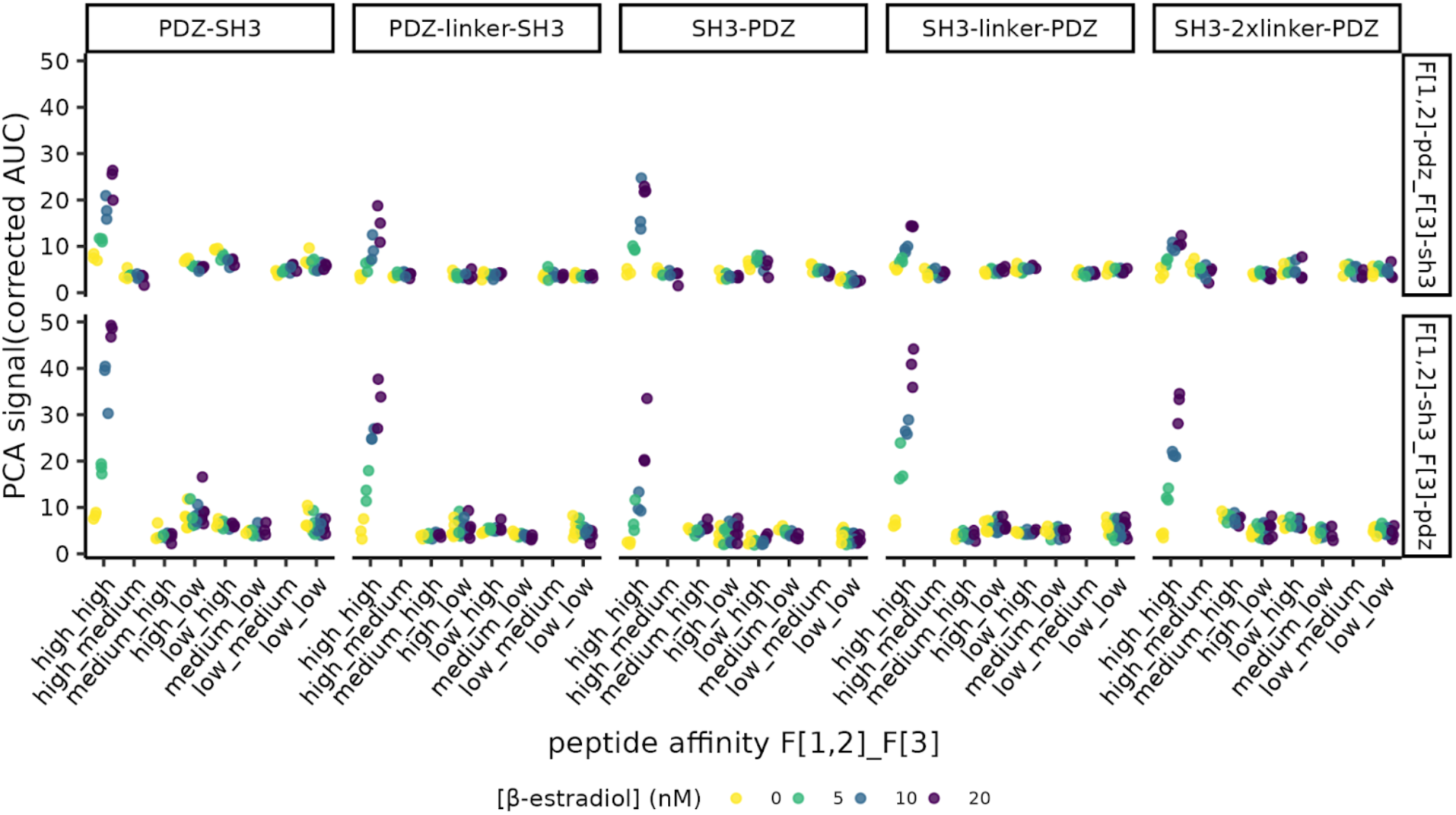
PBD-peptide affinity effect on scaffolding efficiency at different scaffold expression levels. The corrected area under the curve is shown for each scaffold variant, peptide pairs affinity strength and β-estradiol concentration tested in triplicates. The affinity of the PBD-peptide pairs were classified by affinity strength (high : K_d_ < 10 μM, medium : 10 <= K_d_ < 100 μM, low : 100 μM <= K_d_) and the different peptides affinity strength combinations are labelled on the x-axis. The top panels described the peptides in the F[1,2]-pdz.x_F[3]-sh3.x DHFR fragments orientation and the opposite orientation is shown by the bottom panels, i.e F[1,2]-sh3.x_F[3]-pdz.x. The color of the data point indicates the β-estradiol concentration in the PCA media.

**Supplementary Figure 6.**
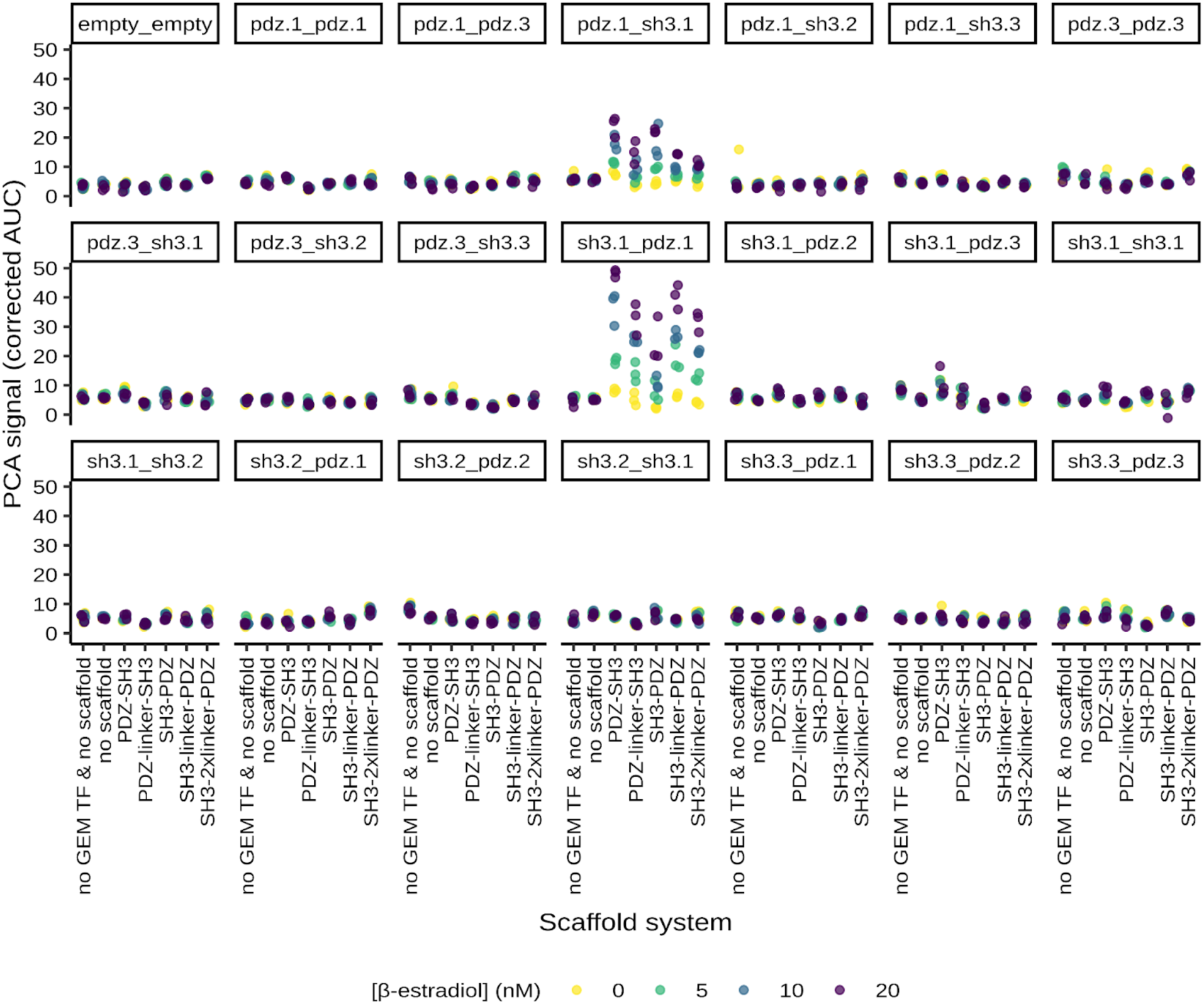
Detailed DHFR PCA results of the scaffolding efficiency assay. The corrected area under the curve is shown for each scaffold system, peptide pairs and β-estradiol concentration tested in triplicates. The peptide combinations are labeled on top of each plot in F[1,2]-peptide_F[3]-peptide orientation. The color of the data point indicates the β-estradiol concentration in the PCA media.

**Supplementary Figure 7.**
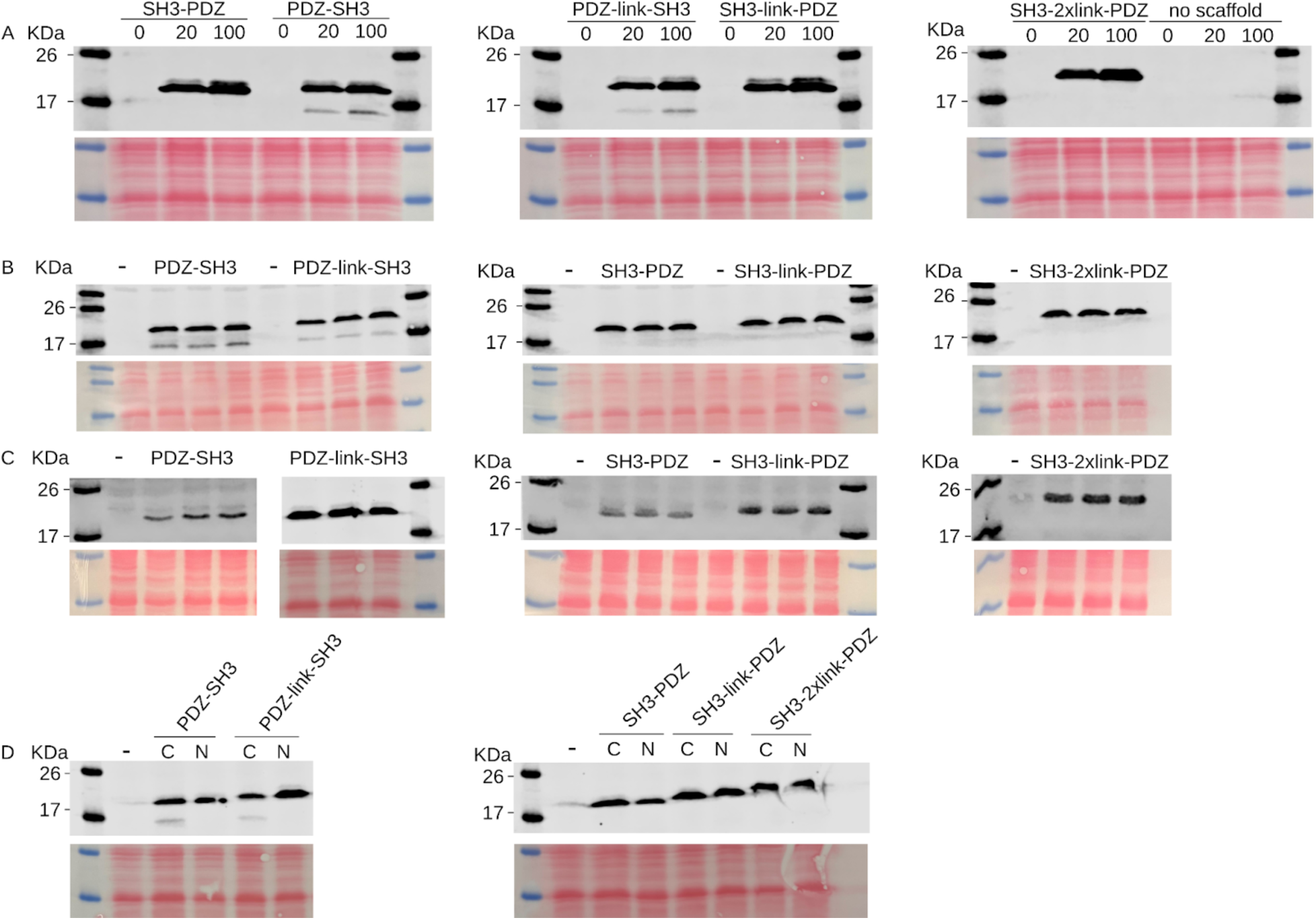
Scaffold induction and abundance validations by Western Blotting. For all panels, the Western Blots are shown above their respective Ponceau coloration, by which we verified the correct transfer of the proteins from the SDS-PAGE gel to the nitrocellulose membrane. The scaffold architectures tagged at the *GAL1* locus with the FLAG epitope (DYKDDDDK) are labelled above each Western Blot. A. Western Blot on protein extracts obtained from yeast strains encoding a C-terminal tagged scaffold grown with 0, 20 or 100 nM β-estradiol. B. Western Blot on protein extracts obtained from yeast strains encoding an untagged (-) or a C-terminal tagged scaffold (n = 3) grown with 20 nM β-estradiol. C. Western Blot on protein extracts obtained from yeast strains encoding an untagged (-) or a N-terminal tagged scaffold (n = 3) grown with 20 nM β-estradiol. FLAG-PDZ-link-SH3 Western Blot was not obtained with the same antibody as the other N-terminal tagged scaffold. D. Western Blot on protein extracts obtained from yeast strains encoding an untagged (-) or a tagged scaffold (N or C-terminal tagged) grown with 20 nM β-estradiol for verification of the tag effect.

**Supplementary Figure 8.**
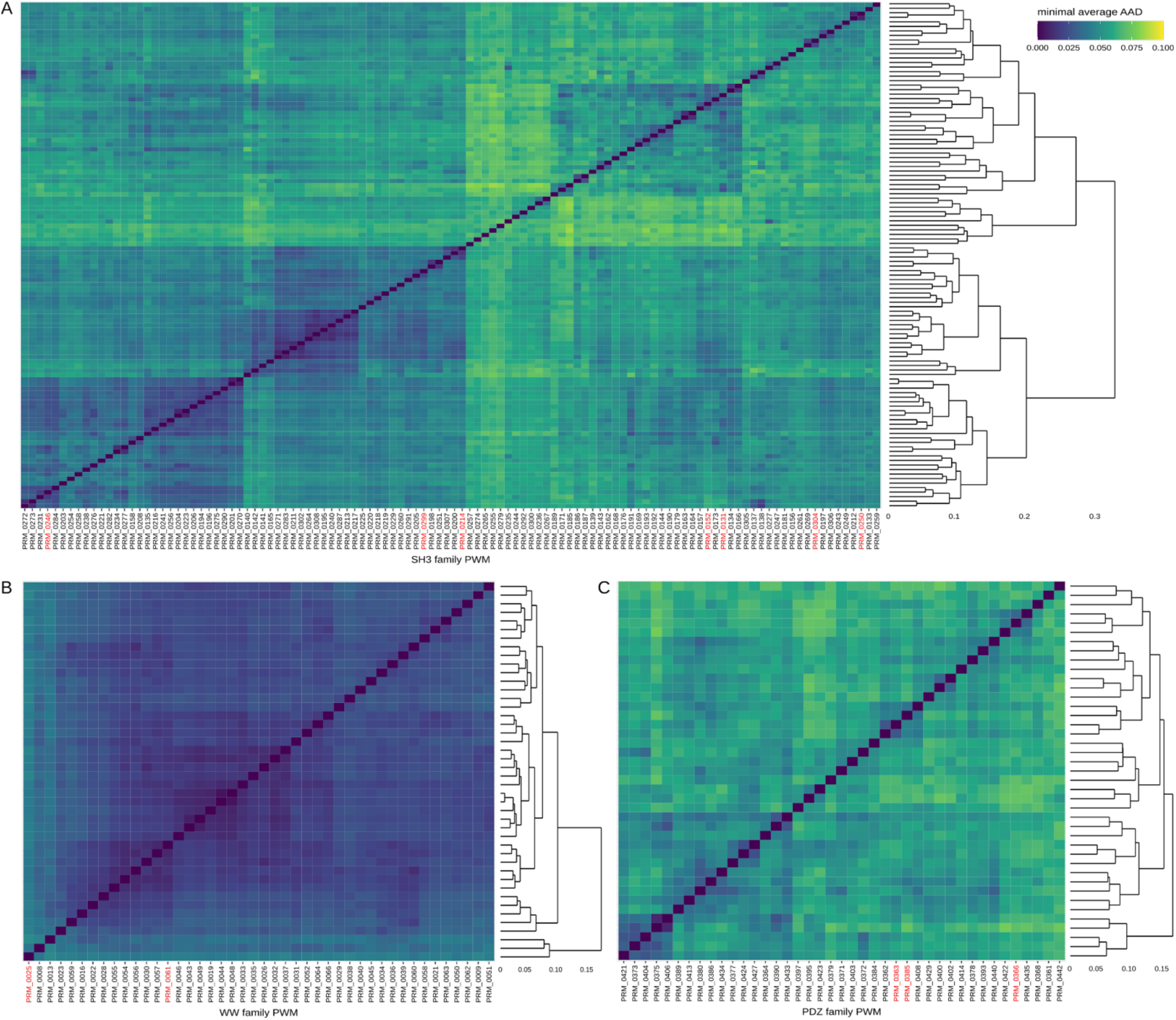
Comparison of PWMs present in the PRM DB^40^. The minimal average AAD computed between each PWMs is represented by the color scale of the heatmaps for the A. SH3, B. WW and C. PDZ family. The dendrograms highlight the clusters of binding specificity. The red labels on the x-axis are the selected PBDs for experiments following visual verification of their PWMs.

**Supplementary Figure 9.**
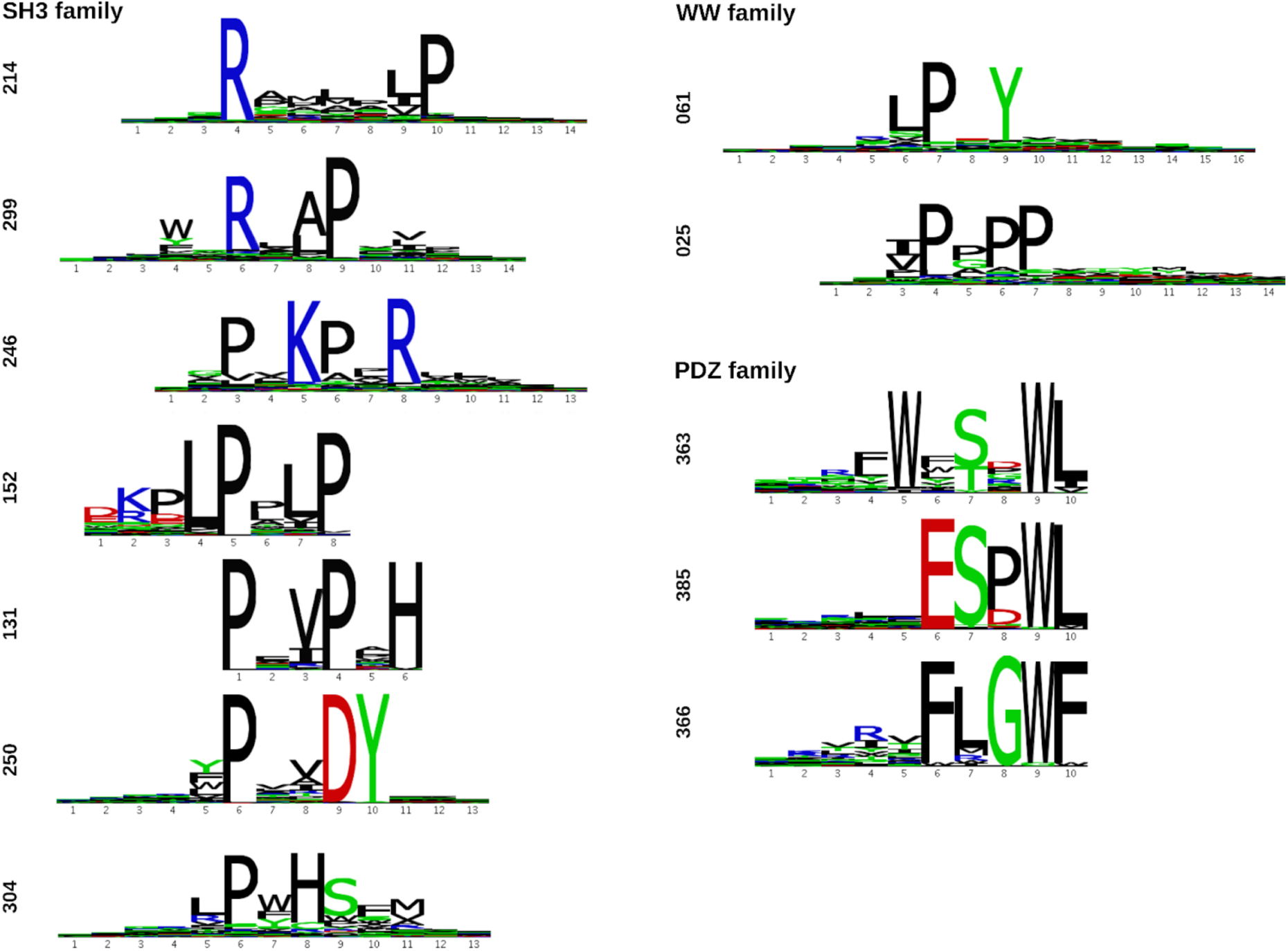
Specificity of the selected PBDs for experiments in yeast. The PWMs extracted from the PRM DB^40^ are aligned to emphasize the conserved binding motif of each PBD family and also the differences in the specificity of each PBD.

**Supplementary Figure 10.**
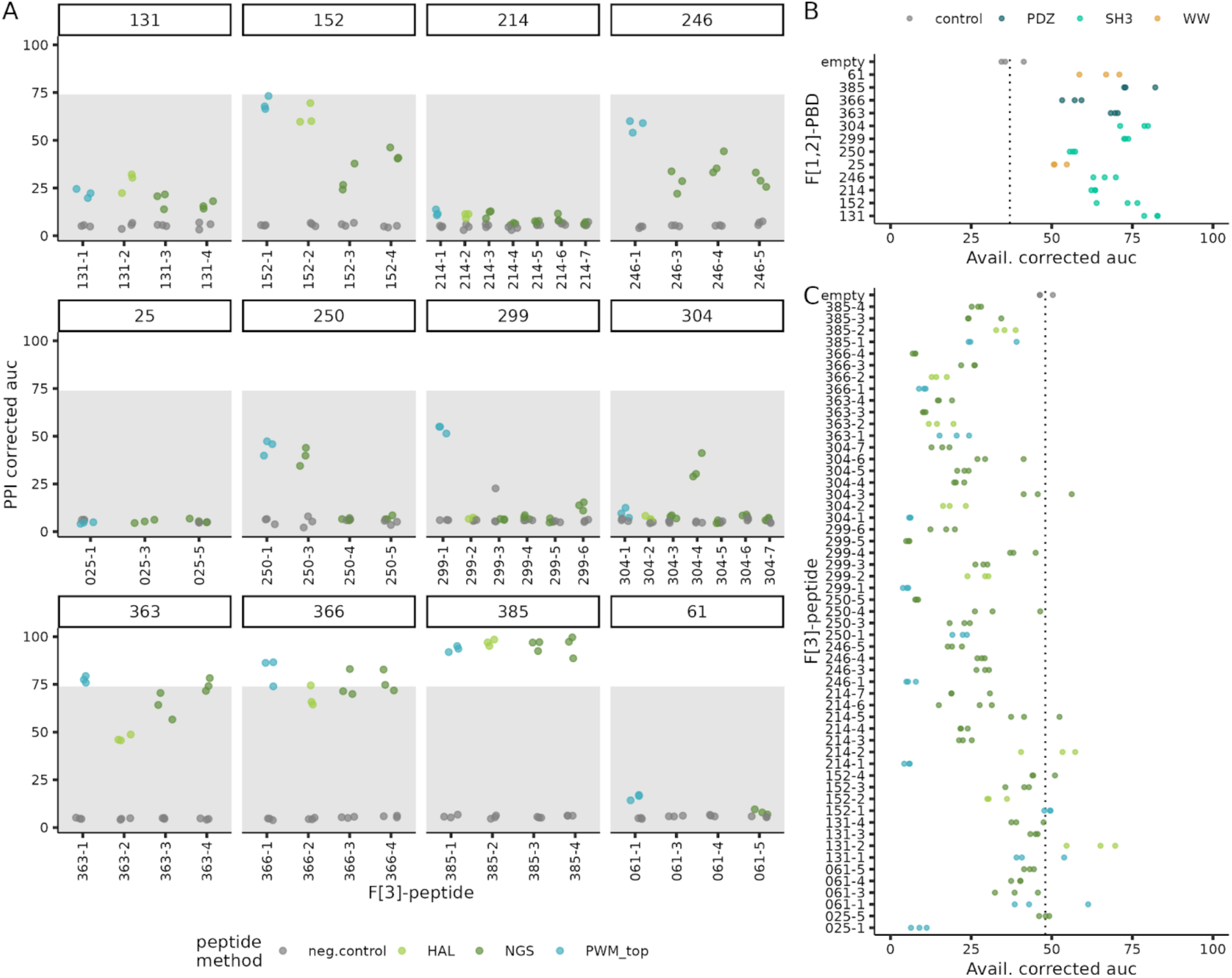
Detailed results for the DHFR PCA experiment with sequences from the PRM DB. A. PCA signal for the PPI strength measurements, in triplicates, is represented for each combination of PBD-peptide. The F[1,2]-PBDs are labelled at the top of each plot and the F[3]-peptides are labelled on the x-axis. The colors of the data points represent the design strategy of each peptide (see methods). The negative controls in grey show the PCA signal obtained when only the F[3]-peptide is present on the plasmid. The shaded area covers the PPI intensity which is not sufficient to be considered as a candidate for scaffold design i.e. PPI score < 12. B. PCA signal obtained for the binding availability of each F[1,2]-PBD in triplicate. The colors of the data points represent the PBD family and the control is the empty plasmid. The vertical dashed line highlights the mean PCA signal of the control. C. PCA signal obtained for the binding availability of each F[3]-peptide in triplicate. The colors of the data points represent the design strategy of each peptide and the control is the empty plasmid. The vertical dashed line highlights the mean PCA signal of the control.

## Notes

https://github.com/Landrylab/Lemieux_et_al2025

### Competing Interest Statement

The authors have declared no competing interest.

### Summary of Updates

Many experiments were added to validate the experimental design, Figure 4 and Supplementary Figures were added. New sequences were characterized.

