## Supplementary File 1 for "A protein-fragment complementation assay to quantify synthetic protein scaffold efficiency"

### Supplementary File 1 - Detailed methods

#### Culture media

The complex medium used for yeast transformation and growth was YPD [1% yeast extract, 2% tryptone, 2% glucose, and 2% agar (for solid medium)] to which we added nourseothricin (NAT, Cedarlane Labs) or hygromycin (HYG, Bioshop Canada) at final concentrations of 100 and 250 µg/ml, respectively. Synthetic medium [SC, 0.174 % yeast nitrogen base without amino acids and without ammonium sulfate, 2% glucose, 0.1% MSG, and 2% agar (for solid medium)] depleted of uracil or leucine (-ura or -leu) was also used for yeast growth and selection. All yeast cultures were incubated at 30°C, with 250 rpm. agitation for liquid cultures. Medium for bacterial growth was 2YT [1 % yeast extract, 1.6% tryptone, 0.2% glucose, 0.5% NaCl and 2% agar (for solid medium)] with ampicillin (AMP) at final concentration of 100 µg/ml. All bacterial cultures were incubated at 37◦C, with 250 rpm agitation for liquid cultures. DHFR PCA selection condition was liquid synthetic media (PCA medium, 0.67% yeast nitrogen base without amino acids and without ammonium sulfate, 2% glucose, drop-out without adenine and uracil) containing 200 µg/ml methotrexate (MTX, BioBasic) diluted in dimethyl sulfoxide (DMSO, Bioshop Canada)

#### PCR mix and cycles

The details on synthetic DNA templates, oligonucleotides and plasmid templates used for all PCR reactions are available in Supplementary Tables 1, 2 and 3, respectively. DNA amplifications were performed with the standard PCR mix and cycle except when mentioned otherwise.

| **Standard PCR mix** | | | | | | |
|---|---|---|---|---|---|---|
| | **Name** | **Concentration** | **Volume (µL)** | **Time** | **Temperature** | **Number of cycles** |
| DNA | Template DNA | 15 ng/µL | 0.75 | 5 min | 98°C | 1x |
| Buffer | Fidelity buffer | 5X | 5 | 20 sec | 98°C | |
| dNTPs | | 10 mM | 0.75 | 30 sec | 60°C | 35x |
| Oligo F | | 10 µM | 0.75 | 1 min / kb | 72°C | |
| Oligo R | | 10 µM | 0.75 | 3 min / kb | 72°C | 1x |
| Enzyme | KAPA | 1 U/µL | 0.5 | | | |
| Water | | | 16.5 | | | |
| | | **Total volume** | 25 | | | |

Colony PCR, for cloning validation, were performed with the following reaction mix:

| Colony PCR mix | | | | | | |
|---|---|---|---|---|---|---|
| | Name | Concentration | Volume (µL) | Time | Temperature | Number of cycles |
| DNA | Cell lysis | 15 ng/µL | 2 | 5 min | 95°C | 1x |
| Buffer | Standard buffer | 5X | 5 | 20 sec | 98°C | 35x |
| dNTPs | | 10 mM | 0.4 | 30 sec | 54°C | |
| Oligo F | | 10 µM | 0.4 | 1 min / kb | 72°C | |
| Oligo R | | 10 µM | 0.4 | 3 min / kb | 72°C | 1x |
| Enzyme | Taq DNA polymerase | 1 U/µL | 0.12 | | | |
| Water | | | 12.68 | | | |
| | | Total volume | 20 | | | |

**Construction of template plasmid**

pKB1 was constructed by cloning the *URA3* marker from pUG72 in the EcoRI/SacI site of M4296[1]. We PCR amplified the *URA3* marker with primer 29-E7 and 29-E8 to add an EcoRI site in 5'. We then double-digested the PCR product with EcoRI and SacI (New England Biolabs, NEB), and purified the digestion on column (QIAGEN) to remove the restriction enzyme. In parallel, we double-digested the M4296 plasmid with EcoRI and SacI (NEB). M4296 double digestion was then dephosphorylated to prevent recircularization of the vector. The resulting dephosphorylated digestion was purified on column (QIAGEN) to remove all enzymes. The double-digested *URA3* marker was cloned in the double-digested M4296 using T4 DNA ligase with a ratio of 3:1 and then transformed into *Escherichia coli* MC1061 chemo-competent cells. Positive clones were confirmed by XhoI (NEB) digestion and Sanger sequencing to confirm proper cloning.

To construct pFA-*mEGFP-NATNT2* plasmid, *mEGFP* gene (insert 1) and *ENO1* terminator fragment (insert 2) were inserted into pFA-*natNT2* (vector) using Gibson assembly[2]. All three fragments were PCR amplified (PCR1, PCR2 and PCR3, respectively) and treated with 20 units of DpnI for 1 h at 37 °C before being purified on magnetic beads. Assembly was performed with the three fragments following a ratio of 1:5:5 (vector:insert 1:insert 2). After transformation in MC1061 strain, a positive clone was retrieved and the integrity of the assembled plasmid was verified by Sanger sequencing.

| PCR1 | | | | | | |
|---|---|---|---|---|---|---|
| | Name | Concentration | Volume (µL) | Time | Temperature | Number of cycles |
| DNA | F65V-34a-mEGFP | 10 ng/µL | 1 | 5 min | 95°C | 1x |

| | | | | | | |
|---|---|---|---|---|---|---|
| Buffer | Fidelity buffer | 5X | 5 | 20 sec | 98°C | |
| dNTPs | | 10 mM | 0.75 | 15 sec | 60°C | 5x |
| Oligo F | 161-F10 | 10 µM | 0.75 | 30 sec | 72°C | |
| Oligo R | 161-F12 | 10 µM | 0.75 | 20 sec | 98°C | |
| Enzyme | KAPA | 1 U/µL | 0.3 | 15 sec | 72°C | 30x |
| Water | | | 16.45 | 30 sec | 72°C | |
| | | **Total volume** | 25 | 3 min | 72°C | 1x |
| | | | | Hold | 10°C | |
| **PCR2** | | | | | | |
| | **Name** | **Concentration** | **Volume (µL)** | **Time** | **Temperature** | **Number of cycles** |
| DNA | pYTK051 | 10 ng/µL | 1 | 5 min | 95°C | 1x |
| Buffer | Fidelity buffer | 5X | 5 | 20 sec | 98°C | |
| dNTPs | | 10 mM | 0.75 | 15 sec | 60°C | 5x |
| Oligo F | 161-G1 | 10 µM | 0.75 | 30 sec | 72°C | |
| Oligo R | 219-B9 | 10 µM | 0.75 | 20 sec | 98°C | |
| Enzyme | KAPA | 1 U/µL | 0.3 | 15 sec | 72°C | 30x |
| Water | | | 16.45 | 30 sec | 72°C | |
| | | **Total volume** | 25 | 3 min | 72°C | 1x |
| | | | | Hold | 10°C | |
| **PCR3** | | | | | | |
| | **Name** | **Concentration** | **Volume (µL)** | **Time** | **Temperature** | **Number of cycles** |
| DNA | pFA-natNT2 | 10 ng/µL | 1 | 5 min | 95°C | 1x |
| Buffer | Fidelity buffer | 5X | 5 | 20 sec | 98°C | |
| dNTPs | | 10 mM | 0.75 | 15 sec | 68°C | 35x |
| Oligo F | 219-A9 | 10 µM | 0.75 | 2 min | 72°C | |
| Oligo R | 161-F9 | 10 µM | 0.75 | 3 min | 72°C | 1x |
| Enzyme | KAPA | 1 U/µL | 0.3 | Hold | 10°C | |
| Water | | | 16.45 | | | |
| | | **Total volume** | 25 | | | |

**Construction of template strain**

Strain AKD0678 was constructed by modifying the *GAL1* locus in strain PL0001. We PCR amplified the DHFR F[1,2]-FLAG-NATMX fragment with primer CLO5-65 and 23-B2 from plasmid pAG25-DHFR[1,2]-linker-FLAG to add stuffer3 from Dionne et al. 2021[3] before the linker-DHFR F[1,2]. We transformed the PCR product in PL0001 using standard lithium acetate yeast transformation[4] to replace the CDS of *GAL1* with the stuffer3. Colony PCR with primer 248-B7 and CLO1-50 and Sanger sequencing confirmed proper fragment integration at *GAL1* locus.

#### Construction of scaffold strains

The peptide binding domains (PBDs, SH3 and PDZ) optimized sequences for expression in *S. cerevisiae* were ordered as gBlock from IDT. They were amplified by PCR with primers adding 40-pb homology arms to the *GAL1* locus in 5' and the other PBD in 3' (PDZ : 312-D8/312-A9, SH3 : 312-A8/312-C9). A second amplification was performed on SH3 and PDZ gBlocks to add 40-pb homology arms to the other PBD in 5' and to the ADHt in 3' (PDZ : 312-B9/312-B8, SH3 : 312-H8/312-E8). Then, a fusion PCR was performed for which a 1:50 dilution of the PCR fragments of the first amplification served as the DNA template. This created the sequences SH3-PDZ (312-D8/312-E8) and PDZ-SH3 (312-A8/312-B8), with 3' homology to the *GAL1* promoter and 5' homology to the ADHt. Also, the SH3 and PDZ sequences were ordered fused with linkers of different lengths (linker : $(GGGGS)_1$-$(GGGAS)_1$, 2xlinker : $(GGGGS)_3$-$(GGGAS)_1$) and position resulting in three additional gBlock (SH3-linker-PDZ, SH3-2xlinker-PDZ, PDZ-linker-SH3). They were amplified using the same primer pairs as the fusion PCRs.

To integrate the different scaffolds at the *GAL1* locus, we added the NATMX selection marker to the gene fragments encoding the scaffolds. We amplified the ADHt-NATMX cassette from the strain AKD0678 to add 40pb-homology arms to either SH3 (312-F8/312-G8) or PDZ (312-C8/312-G8) sequence (PCR4). Then, we performed a fusion PCR with the fragments encoding the scaffolds and the ADHt-NATMX cassettes (SH3-x-PDZ : 312-A8/312-G8, PDZ-x-SH3:312-B8/ 312-G8). The resulting gene fragments, scaffold-ADHt-NATMX, were transformed by homologous recombination in PL0001 using standard lithium acetate yeast transformation [4] and plated on selective media (YPD+NAT), resulting in the strains PL0012 to PL0016 (Supplementary Table 5). The correct genomic integrations were validated by colony PCR (248-B7/CLO1-50) and Sanger sequencing.

| PCR 4 | | | | | | |
|---|---|---|---|---|---|---|
| | **Name** | **Concentration** | **Volume (µL)** | **Time** | **Temperature** | **Number of cycles** |
| DNA | AKD0678 genomic DNA | 50 ng/µL | 2 | 5 min | 98°C | 1x |
| Buffer | Fidelity buffer | 5X | 5 | 20 sec | 98°C | 35x |
| dNTPs | | 10 mM | 0.75 | 30 sec | 60°C | |

| | | | | | | |
|---|---|---|---|---|---|---|
| Oligo F | | 10 µM | 0.75 | 1 min / kb | 72°C | |
| Oligo R | | 10 µM | 0.75 | 3 min / kb | 72°C | 1x |
| Enzyme | KAPA | 1 U/µL | 0.5 | | | |
| Water | | | 15.25 | | | |
| | | **Total volume** | 25 | | | |

The scaffold sequences were inserted again at the *GAL1* locus fused with a FLAG epitope (DYKDDDDK) in N-terminal or in C-terminal. The same PCR fusion strategy was used to construct the FLAG-tagged scaffold by replacing the primers in 5' or 3' extremities of the scaffold sequence with the ones encoding the FLAG epitope (Supplementary Table 2) for the first PCR (standard PCR mix and cycle). The second PCR for the FLAG C-terminal tagged scaffolds used the same primers as the untagged scaffolds, but the FLAG N-terminal tagged scaffold second PCR used a different primer pair (360-D12/312-G8). The resulting gene fragments were transformed and validated with the same approach as the strains PL001X, resulting in 10 strains encoding at their *GAL1* locus scaffolds fused to a FLAG epitope (N-terminal FLAG : PL0039 to PL0043, C-terminal FLAG : PL0023 to PL0027, Supplementary Table 5).

**Validation of β-estradiol induction**

In order to validate the expression range of the β-estradiol inducible system, we constructed a strain expressing a fluorescent protein, mEGFP, from the *GAL1* locus in the PL0001 background strain. The mEGFP-NATNT2 gene fragment was then amplified from the purified pFA-mEGFP-NATNT2 plasmid with the standard PCR mix adding 40-pb homology to the *GAL1* locus and a linker sequence in N-terminal of the mEGFP (CLO5-61/61-G8). The resulting gene fragment was transformed by homologous recombination in PL0001 using standard lithium acetate yeast transformation[4] and plated on selective media (YPD+NAT), resulting in the strain AKD0840 (Supplementary Table 5).

The strain AKD0840 was grown in SC-leu overnight in triplicate, then was diluted at 0.15 OD in SC-leu with different concentrations of β-estradiol, ranging from 0 nM to 50 nM. The cultures with β-estradiol were grown until the exponential phase was reached (OD = 0.5 - 0.7). Cultures were then diluted to 0.05 OD in sterile water for cytometry measurements. The fluorescent intensity of 5000 cells per replicates was measured by a Guava EasyCyte HT cytometer (blue laser, λ = 488 nm). The median fluorescent intensity of each replicate is computed and used to follow the β-estradiol concentration effect on induction.

**Protein abundance assay by Western blotting**

To validate the proper expression of the scaffold with the β-estradiol induction, we used the strains PL0023 to PL0027 encoding the 5 scaffold architectures tagged with a FLAG epitope in

C-terminal. The strains were grown overnight in YPD. The next day the strains were diluted at 0.15 OD in SC-complete with β-estradiol (0 nM, 20 nM and 100 nM) and grew up to 1 OD. The cell pellets were collected and kept at -80$_{\circ}$C for further use. The pellets were resuspended at 0.05 OD/µL in lysis buffer (cOmplete, Mini Protease Inhibitor Cocktail, Roche) and 250 µL was used for cell lysis. Glass beads were added to the cell suspension and proceeded to vortex the mixture in a Turbomix for 5 minutes. Then, 25 µL of 10% SDS was added to the samples and they were boiled for 10 minutes. The samples were centrifuged at 16,000g for 5 minutes and the supernatant was collected. Samples for migration were prepared by mixing 17.5 µL of clear supernatant, 2.5 µL of 1M DTT and 5 µL of 5X loading buffer (250 mM Tris-Cl pH 6.8, 10% SDS, 0.5% bromophenol blue, 50% glycerol) and boiled for 5 minutes. Samples were migrated on a 15% SDS-PAGE gel and transferred on a nitrocellulose membrane at 0.8 mA/cm$^2$ for 1h15min. Each membrane was stained with Ponceau to confirm proper loading and appropriate transfer. The membrane was blocked overnight in a blocking buffer (Intercept® (PBS) Blocking Buffer). The next morning, an Anti-FLAG antibody (1/10,000, Monoclonal ANTI-FLAG® M2, Sigma-Aldrich or THE Anti Flag-Tag, GenScript) was incubated with the membrane for 2 hours in the blocking buffer with 0.2 % Tween 20. The membrane was then washed with PBS-T (100mM NaCl, 80mM $Na_2HPO_4$, 25mM $NaH_2PO_4$, 0.1% Tween 20) five times and incubated 1 hour with the secondary antibody (1/10,000, Anti-Mouse 800, Mandel Scientific) in blocking buffer added with 0.2% Tween 20. The membrane is then washed with PBS-T 5 times and imaged with an Odyssey Fc instrument (Licor, Mandel Scientific) in 700 and 800 channels.

The protein extraction and western blot protocol were repeated for cell pellets collected in triplicate with 20 nM β-estradiol for the strains PL0023 to PL0027, PL0039 to PL0043 and PL0012 to PL0016 (used as negative controls).

**PBD and peptide sequence selection from the PRM DB**

PCR cycle used to amplify peptide sequences to insert in frame F[3]-linker.

| **PCR5** | | | | | | |
|---|---|---|---|---|---|---|
| | **Name** | **Concentration** | **Volume (µL)** | **Time** | **Temperature** | **Number of cycles** |
| DNA | pGD110 | 10 ng/µL | 0.75 | 5 min | 95°C | 1x |
| Buffer | Fidelity buffer | 5X | 5 | 20 sec | 98°C | |
| dNTPs | | 10 mM | 0.75 | 15 sec | 42°C | 5x |
| Oligo F | A | 10 µM | 0.75 | 10 sec | 72°C | |
| Oligo R | 312-F4 | 10 µM | 0.75 | 20 sec | 95°C | |
| Enzyme | KAPA | 1 U/µL | 0.5 | 15 sec | 60°C | 17x |
| Water | | | 16.5 | 10 sec | 72°C | |
| | | **Total volume** | 25 | 1 min | 72°C | 1x |
| | | | | Hold | 10°C | |

A : 344-A4 to 344-H10

**Construction of pPL9-variants**

To construct pPL9, we used the pGD110 backbone and removed the DHFR fragments to replace them by an mEGFP. First, pGD110 was digested with BamHI-HF enzyme and purified with magnetic beads. The linear plasmid was then used as template DNA for a mutagenesis of the backbone and removing the pCYC-DHFR F[1,2] using the standard PCR (305-G8/H8). The resulting linear fragment was transformed in MC1061 chemo-competent cells and the cells were plated on selective media (2YT+AMP). The pGD110-pCYC-F[3]-CYCt (pPL8) was validated by enzymatic digestion (BamHI-HF and HindIII-HF) and by whole-plasmid sequencing (Plasmidsaurus). pPL8 was then double digested with HindIII-HF and XbaI, to remove F[3], and migrated on an 1.2% agarose gel to purify the larger fragment. Then, the mEGFP sequence was amplified from pFA-mEGFP-NATNT2 (PCR6) and the linker-CYCt was amplified from pGD110 (Standard PCR, 352-C1/D1), both with 40-pb homology to the designed neighbor fragments. The two gene fragments were purified with magnetic beads and used as inserts in a Gibson assembly with the double digested pPL8 backbone using a 3:1:1 ratio (backbone : insert 1: insert 2)[2]. After transformation in *Escherichia coli* MC1061 strain, clones were validated by colony PCR and the purified plasmid (pPL9) was sent to whole-plasmid sequencing (Plasmidsaurus).

| PCR6 | | | | | | |
|---|---|---|---|---|---|---|
| | **Name** | **Concentration** | **Volume (µL)** | **Time** | **Temperature** | **Number of cycles** |
| DNA | pFA-mEGFP-NATNT2 | 15 ng/µL | 1 | 5 min | 95°C | 1x |
| Buffer | Fidelity buffer | 5X | 5 | 20 sec | 98°C | 35x |
| dNTPs | | 10 mM | 0.75 | 15 sec | 52°C | |
| Oligo F | 352-A1 | 10 µM | 0.75 | 1 min | 72°C | |
| Oligo R | 352-B1 | 10 µM | 0.75 | 3 min | 72°C | 1x |
| Enzyme | KAPA | 1 U/µL | 0.3 | Hold | 10°C | |
| Water | | | 16.45 | | | |
| | | **Total volume** | 25 | | | |

To insert peptide sequences in frame with the mEGFP-linker present on pPL9, we used the same cloning approach as for pGD110-variants since the insertion site is the same in both plasmids. The pPL9-variants were sequenced by Sanger sequencing (CHUL sequencing platform).
